# ChAI-Net learns a regulatory code for chromatin co-accessibility

**DOI:** 10.64898/2026.10.29.762938

**Authors:** Jodi Lee, Kaite Zhang, Ethan Shurberg, Vijay Ramani, Katherine S. Pollard

## Abstract

The regulation of chromatin accessibility is critical for cell function and identity. Advances in single-molecule long-read footprinting have enabled near-nucleotide-resolution readout of chromatin fiber organization, capturing both coordination of transcription factor (TF) binding and variation in nucleosome positioning previously obscured by bulk methods. Here we combine this technique with deep learning to study how protein-DNA interactions are coordinated within different types of *cis*-regulatory elements in mouse embryonic stem cells. First, we quantify the frequency with which pairs of positions are simultaneously accessible on individual molecules. Then, we train ChAI-Net, a neural network that predicts these locus-specific chromatin co-accessibility maps from the underlying DNA sequence. Through model interpretation and *in silico* experiments, we show that ChAI-Net learns unique combinations of sequence motifs underlying co-accessibility within enhancers, promoters, and CTCF-bound sites. Many of these motifs are missed when modeling average accessibility, suggesting a specific role in coordinated binding. ChAI-Net’s regulatory grammar also highlights the importance of TF motif orientation, spacing, positioning, and co-occurrence in driving co-accessibility. Thus, our framework for extracting regulatory rules from footprinting data expands understanding of TF cooperativity, while laying the groundwork to elucidate how these rules vary across cellular contexts as single-molecule genomics is deployed at scale.

## Introduction

Chromatin accessibility is a defining feature of eukaryotic genome regulation, describing the physical access of nuclear molecules to chromatinized DNA^1^. Genome-wide profiling of chromatin accessibility has led to the identification of *cis*-regulatory elements (CREs), non-coding DNA sequences harboring clusters of transcription factor (TF) binding sites^2^. Increasing evidence suggests that the combinatorial action of multiple TFs likely shapes the *cis-*regulatory code for TF binding, CRE activity, and regulation of gene expression^3–5^. While these studies established TF cooperativity as a prevalent mechanism of regulating CREs, the use of conventional genomic assays (i.e. ATAC-seq^6^ , DNase-seq^7^, and MNase-seq^8^), which produce average measurements across thousands of cells, hinders our ability to measure events co-occurring on the same DNA molecule^9^.

To overcome these limitations, single-molecule genomic footprinting methods^10–16^ resolve the molecular states of individual DNA molecules at nucleotide resolution across hundreds to thousands of base pairs, revealing the distribution and coordination of accessible sites across individual chromatin fibers^9^. This has enabled the detection of cooperative TF occupancy on individual DNA molecules^3,17,18^, as well as coordinated activity between distal regulatory elements along the same chromatin fiber. As a result, direct measurement of the frequency with which genomic sites are co-accessible across different reads aligning to the same locus could reveal regulatory dependencies between CREs, including cooperative or antagonistic relationships.

Despite these methods, deciphering the *cis*-regulatory code remains a challenge. While sequence-to-function neural networks have been key to learning *cis*-regulatory sequence rules from DNA sequence^2^, most models have been trained on averaged measurements of molecular readouts^19–22^, limiting the discovery of coordinated binding on the same DNA molecule^2^.

Here, we leverage data generated in mouse embryonic stem cells (mESCs) using the **s**ingle-molecule **a**denine **m**ethylated **o**ligonucleosome **s**equencing **a**ssay (SAMOSA)^16,23,24^ to train a neural network we term **Ch**romatin co**A**ccessibility **I**dentifier **Net**work (ChAI-Net). ChAI-Net is designed to learn the DNA sequence elements contributing to co-accessibility, the frequency at which two genomic sites are simultaneously accessible along the same molecule. We hypothesized that modeling co-accessibility will capture a *cis-*regulatory code that reflects cooperative and competitive TF binding patterns. Such a model would enable the inference of the functional relationships between genomic sites.

We find that ChAI-Net learns an expanded set of discovered (*de novo*) motifs compared to modeling average DNA accessibility, and these motifs differ across enhancers, promoters, and CTCF-bound regulatory elements. ChAI-Net *de novo* motif instances show selective co-occurrence and preferred binding positions within regulatory elements. We identified TF motif pairs with recurrent spacing, such as NFY and CTCF separated by ∼19 base pairs (bp), and a dependency on CTCF motif orientation. Furthermore, *de novo* motifs identified from promoters of expressed genes displayed stereospecific alignment on the same face of the DNA helix, suggestive of TF cooperativity. Thus, ChAI-Net offers a new approach to study the *cis-*regulatory code mediating coordinated accessibility across kilobase pair lengths, revealing sequence motifs contributing to coordinated TF binding at CREs.

## Results

### SAMOSA captures chromatin co-accessibility across kilobase pair distances

SAMOSA provides near-bp resolution of protein-DNA interactions across kilobase (kb) distances. We hypothesized that kb-scale accessibility patterns along individual fibers may reveal the sequence motifs of synergistic and antagonistic binders and that variation in these patterns across fibers aligned to a given locus could be used to learn the frequency of coordinated accessibility (**Fig. 1a**). To test our hypothesis, we leveraged a total of ∼10.5 million individual SAMOSA reads with mean length 6,096 bp from mESCs^23^ (E14 line; **Extended Data Fig. 1**). Mean genome-wide sequencing depth was 0.48x, with a 36.7% of the genome covered equal to or greater than 1x and 8.8% covered equal to or greater than 2x (**Methods**).

**Figure 1.**
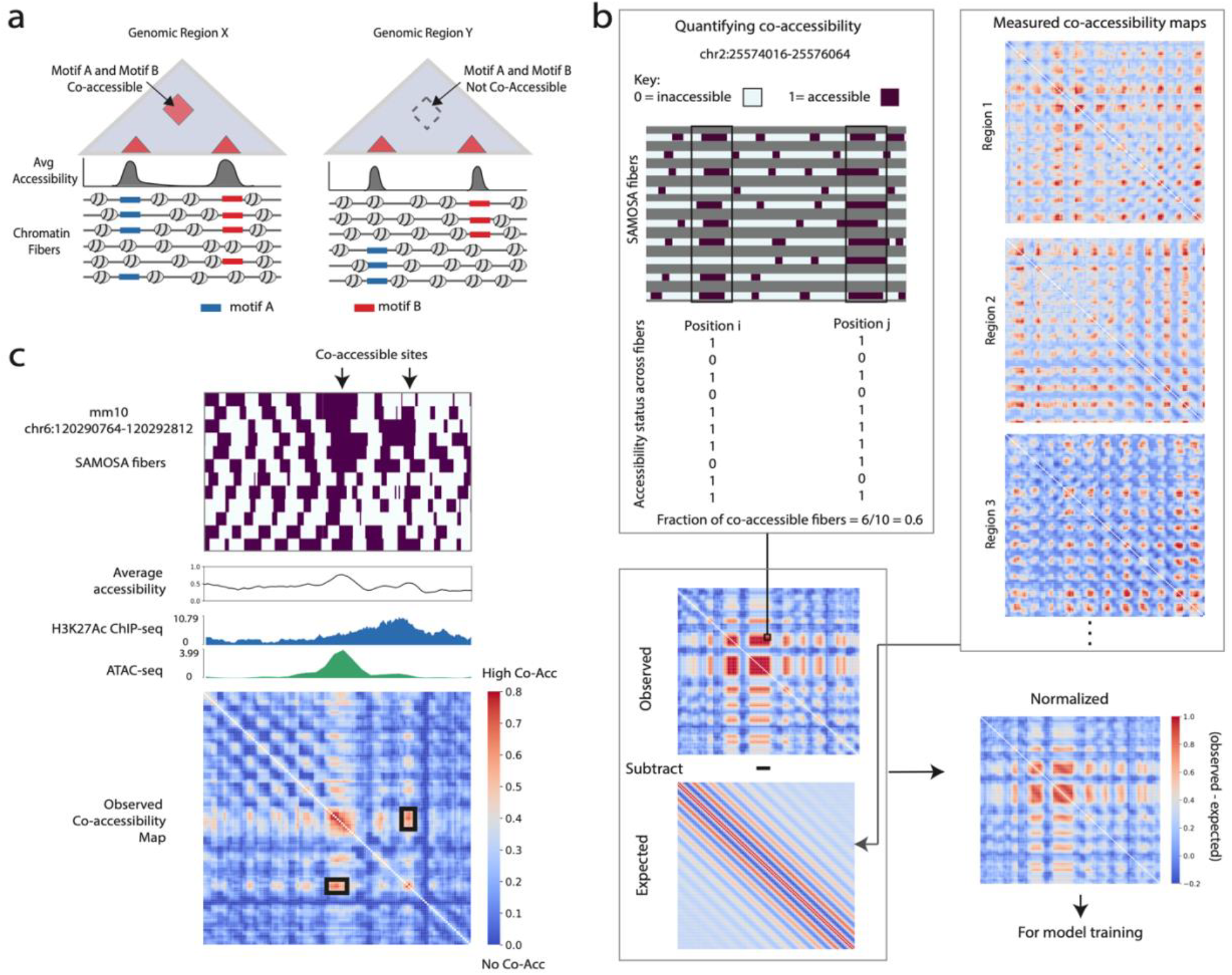
Co-accessibility maps capture pairs of simultaneously open chromatin sites. **a**. Chromatin co-accessibility (left) of sequence elements, such as pairs of TF binding sites, can be differentiated from accessible but uncoordinated elements (right) using single-molecule footprinting. **b**. Observed co-accessibility is computed in a 2,048-bp window using binary accessible vs. inaccessible calls on aligned SAMOSA sequencing reads. This matrix is normalized by subtracting the expected co-accessibility matrix (estimated the average over observed maps). **c**. Example of a window (mm10 chr6:120290764-120292812) with pairs of binds showing higher than expected co-accessibility (red) alongside the underlying SAMOSA chromatin fibers, chromatin accessibility (average over SAMOSA fibers, ATAC-seq signal), and H3K27Ac ChIP-seq. Black box marks a co-accessible event.

In order to stratify our explorations by genomic context, we used publicly available mESC data to annotate regulatory elements genome-wide (**Methods**), including expressed transcription start sites (TSSs), enhancers, and binding sites of CTCF, a key regulator of chromatin interactions with various co-factors that vary across cell types^25^. Specifically, we classified TSSs as expressed (transcripts per million (TPM) > 0) or not using gene expression data from the Mouse ENCODE Project^26–28^, we defined enhancers as promoter-distal ATAC-seq peaks with associated PRO-seq signal^29^, and we annotated CTCF binding sites as ChIP-seq peaks with high Find Individual Motif Occurrences (FIMO) scores^30,31^.

To quantify the variation in accessibility across aligned fibers genome wide, we split the mouse genome (mm10) into 2048 bp genomic windows and calculated a co-accessibility map for each window by computing the frequency of co-occurring accessible sites across footprinted fibers for each pair of 16 bp bins within the window (**Fig. 1b**). We observed that many of the resulting co-accessibility maps had grids of high co-accessibility dots, reflecting periodic open chromatin between uniformly spaced nucleosomes (*i*.*e*. accessible ‘linker’ DNA). The intensity of these dots tended to fade with distance, consistent with nucleosome spacing being most similar between adjacent nucleosomes. Even though this grid-like pattern is consistent genome-wide, the mean across co-accessibility maps has periodic diagonal bands, reflecting the random positioning of windows relative to the first nucleosome as well as variability in fiber types with different nucleosome repeat lengths^12,16^.

Some regions, including those containing enhancers, showed regions of co-accessibility along the same molecule among a subset of aligned SAMOSA fibers and were detected by the co-accessibility measurement as heat off the diagonal (**Fig. 1c**). This is a regulatory event uniquely detectable using single-molecule co-accessibility, because co-occurring, co-excluding, and randomly occurring accessibility look the same when averaging accessibility over SAMOSA fibers, as they do in ATAC-seq data.

Next, to explore sequence features that alter co-accessibility signals relative to the genome-wide expectation, we subtracted the genome-wide average co-accessibility map from each observed map, creating normalized matrices (**Fig. 1b**). To avoid leaky pre-processing when training sequence-based deep learning models, this normalization was performed separately for pre-defined training, validation, and test sets representing an 80/10/10 cross-validation setup^32^.

### ChAI-Net predicts chromatin co-accessibility patterns from DNA sequence

To identify *cis-*regulatory sequence features predictive of chromatin co-accessibility, we developed a deep convolutional neural network (CNN), called ChAI-Net (**Fig. 2a, Methods**). Briefly, ChAI-Net extracts genomic sequence features and learns their combinations, such that the learned representation can reconstruct the normalized co-accessibility map. The model takes 2048 bp of DNA as input and predicts the corresponding 2D map at 16-bp resolution. We chose the ∼2-kb length scale to focus on coordinated binding of TFs and nucleosomes within regulatory elements. This window size was also optimal in terms of being as large as possible while ensuring enough SAMOSA footprinted fibers to estimate co-accessibility of 16-bp bin pairs on the same molecule.

**Figure 2.**
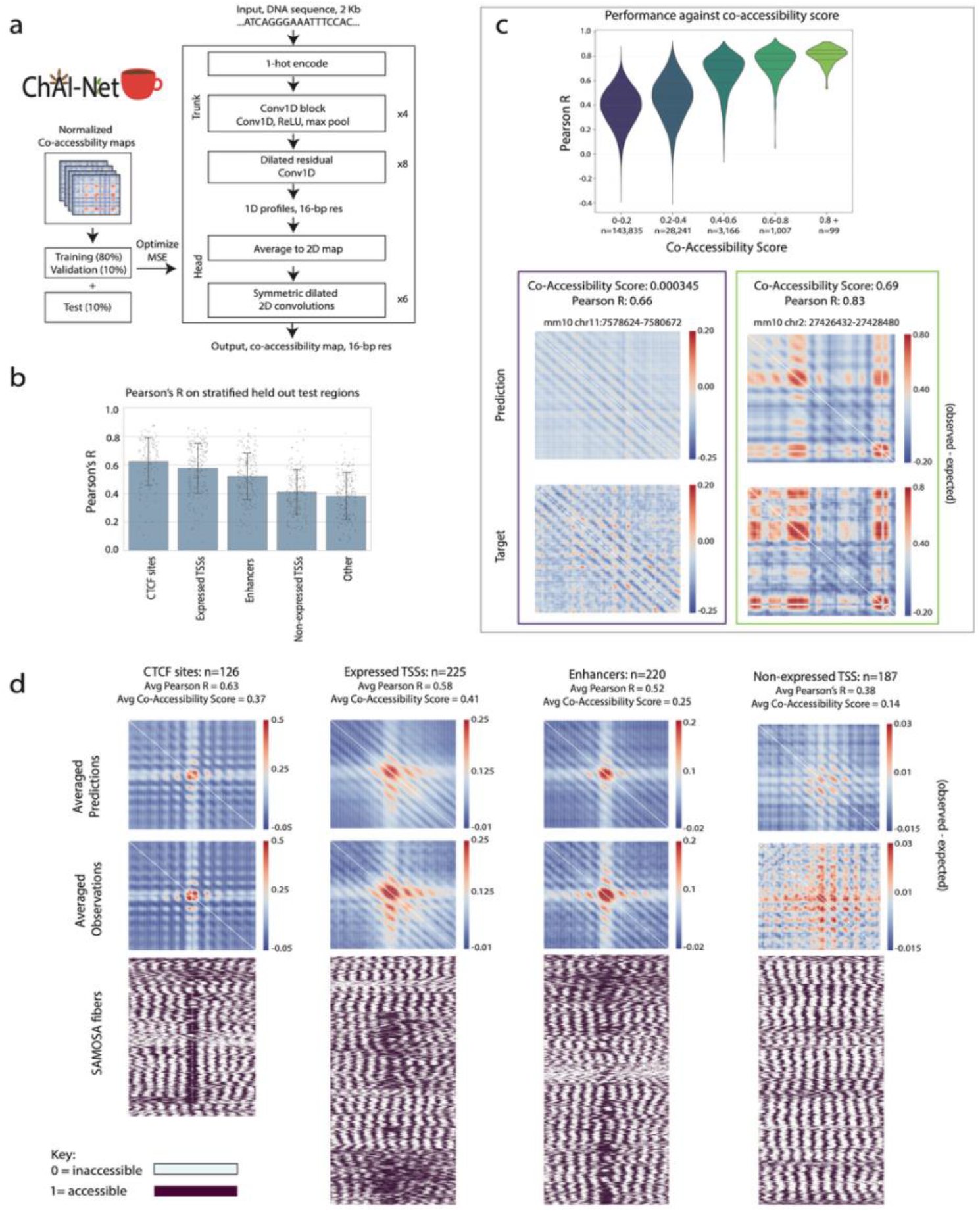
ChAI-Net: a neural network for predicting co-accessibility patterns from DNA sequence. **a**. The ChAI-Net architecture and training strategy (details in **Methods**). **b**. Held-out test set performance as Pearson’s *r* stratified by overlapping regulatory regions (CTCF sites, expressed TSSs, enhancers, non-expressed TSSs) or not. **c**. Distribution of *r* over test windows binned by amount of observed co-accessibility (top) and examples of predicted and observed co-accessibility maps (below) from a window with low co-accessibility (dark purple box) and a window with high co-accessibility (lime green box). **d**. Predicted and observed co-accessibility map averages over test windows overlapping each type of regulatory element. Shown below are the SAMOSA fibers from all windows contributing to each average.

ChAI-Net is trained from scratch on ∼3.4 million genomic windows, but its architecture is an extension of Akita, a previous CNN model that accurately predicts 3D chromatin interaction maps derived from bulk measurements (e.g., Hi-C)^33^. Key modifications included changing the window size and resolution and training on co-accessibility rather than interaction frequency (**Methods**). During training, we used the mean squared error (MSE) between predicted and observed maps as the loss function. To evaluate performance on the held-out validation and test sets, we used both MSE and Pearson’s correlation (*r*). We explored modifications of the hyper-parameters batch size and learning rate, using the validation set to select a parameter combination that maximized performance (**Methods**).

The resulting ChAI-Net model learned a predictive representation of chromatin co-accessibility from DNA sequence (0.0079 MSE, 0.46 *r* on the test set). CTCF sites, enhancers, and TSSs showed better performance than other genomic windows, with expressed TSSs outperforming non-expressed ones (**Fig. 2b, Methods**). These trends suggest a stronger relationship between sequence and co-accessibility in active regulatory elements. More broadly, regions with higher observed co-accessibility were better predicted by ChAI-Net (median *r >* 0.8 for windows with the most co-accessibility versus ∼0.4 for the least co-accessible windows; **Fig. 2c**). Furthermore, ChAI-Net correctly predicted co-accessible patterns that were unique to each type of regulatory element. These included known phasing of nucleosomes flanking CTCF motifs, +1 nucleosomes positioned at expressed TSSs versus low co-accessibility at inactive TSSs, and nucleosome-free regions with well-positioned flanking nucleosomes at enhancers (**Fig. 2d**). These results indicate that ChAI-Net effectively learned relationships between co-accessibility patterns and the underlying DNA sequences.

### Accessibility and co-accessibility are governed by partially distinct sequence motifs

Next, we leveraged a series of model interpretation approaches to investigate what sequences ChAI-Net uses to predict the co-accessibility maps of 23,839 enhancers, 20,026 expressed TSSs, and 11,097 CTCF sites (**Fig. 3a, Methods**). For each regulatory element centered window, we first quantified the contribution of each base to predicting co-accessibility. Next, we aggregated contribution scores into *de novo* motifs, represented as contribution weight matrices (CWMs) in which the magnitude of the weight for a given base quantifies its importance to the predictions and its sign indicates a positive versus negative effect^37,38^. The resulting *de novo* motifs were annotated using similarity to TF position weight matrices (PWMs) from JASPAR and HOCOMOCO databases^34,35^. Finally, instances of sequences matching each CWM were identified within every regulatory element centered window.

**Figure 3.**
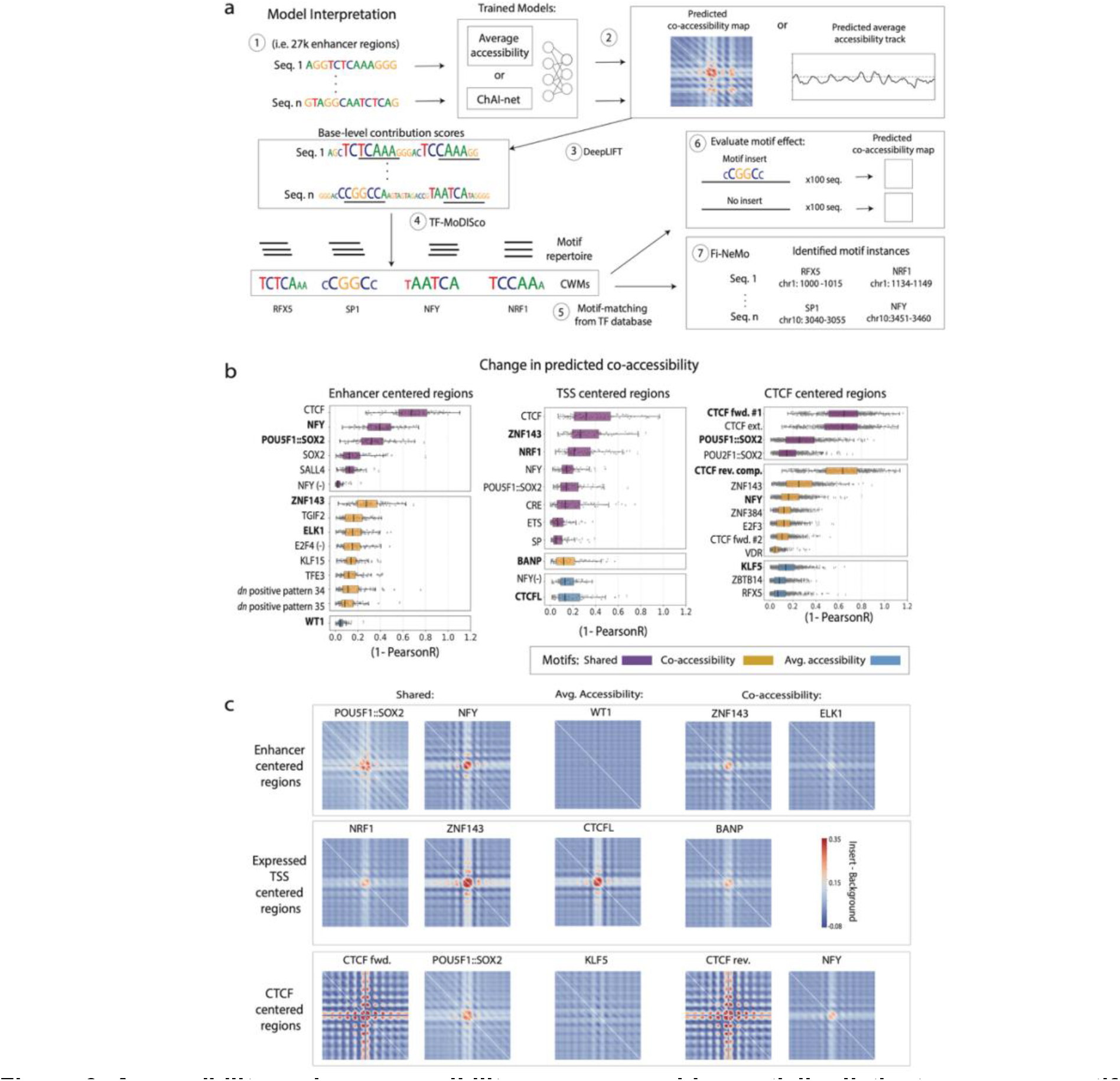
Accessibility and co-accessibility are governed by partially distinct sequence motifs. **a**. Model interpretation workflow. Each 2,048-bp DNA sequence centered on an enhancer, expressed TSS, or CTCF site was used to predict a 2D co-accessibility map with ChAI-Net and a 1D accessibility track with the average accessibility model. Predictions were analyzed by DeepLIFT^34^ to derive per-base scores representing contribution to the prediction (co-accessibility or average accessibility), and scores were combined into contribution weight matrices (CWMs) that represent contribution patterns shared across loci within each type of regulatory region. CWMs were then annotated using TF motif databases (JASPAR^34^, HOCOMOCO^35^). Finally, specific instances of subsequences matching each CWM were identified within each DNA sequence using Fi-NeMo^36^ (**Methods**). **b**. Amount of predicted change (1-PearsonR) in co-accessibility as a result of inserting the consensus sequence of a given CWM into 100 background DNA sequences (**Methods**). CWMs are labeled if a significant PWM-CWM match was found, and they are stratified by whether they were discovered in enhancers (left), expressed TSSs (center) or CTCF sites (right) and colored to indicate if they were discovered by ChAI-Net (yellow), the average accessibility model (blue) or both (purple). Bolded motif names correspond to the maps shown in (**c**). **c**. Change in predicted co-accessibility: average over 100 predicted maps with the motif inserted in the center minus average over 100 predicted maps without the motif. Diagonal values are not plotted to emphasize bin x bin co-accessibility (white).

ChAI-Net motifs varied across the three types of regulatory elements (**Extended Data Fig. 2, Extended Data Fig. 3**). For example, SALL4, ELK1, KLF15, TFE3, and TGIF2 were important in enhancers only, whereas expressed TSSs uniquely highlighted NRF1, BANP, CRE, ETS, and SP, while CTCF sites uniquely highlighted ZNF384 and VDR. TF motifs learned from all three types of regulatory elements included NFY, CTCF, POU5F1::SOX2, and ZNF143. These findings suggest that co-accessibility occurs at the binding sites of TFs that are both ubiquitous across regulatory contexts and those that bind specifically at one type of regulatory element.

We reasoned that if motifs that specifically contribute to co-accessibility exist, they would be distinct from those that contribute to one-dimensional (1D) accessibility. To test this hypothesis, we trained a separate model that uses the trunk of ChAI-Net to predict average accessibility of SAMOSA fibers (**Extended Data Fig. 4**) and applied the same motif discovery pipeline to the same regulatory regions using the average accessibility model. We then compared the discovered motifs between the two models (**Methods**). TF matching motifs were classified as being derived from ChAI-Net, the average accessibility model, or both.

Both models used a relatively small set of *de novo* motifs for making predictions (range: 9 to 35 per regulatory element type). However, compared to the average accessibility model, ChAI-Net discovered more motifs matching known TF binding preferences, and its motif set also differed notably from those of the average accessibility model (**Extended Data Fig. 2, Extended Data Fig. 5**). TFs uniquely highlighted by ChAI-Net include VDR, TGIF2, TFE3, ZNF384^39,40^, and CTCF’s reverse orientation motif. ChAI-Net placing importance on CTCF’s reverse complement motif suggests that modeling co-accessibility may have required learning patterns that depend on CTCF motif orientation, which we explore further below. The average accessibility model uniquely highlighted several TFs, including WT1, which guides formative pluripotency progression and whose forced expression leads to the transition of naïve pluripotent cells towards a post-implantation epiblast identity^41^.

While CWMs quantify the importance of motifs to predicting co-accessibility, they do not establish if TFs are sufficient to drive coordinated binding. To test this computationally, we evaluated each learned motif’s ability to change predicted co-accessibility by inserting its consensus sequence into 100 background sequences matched for GC-content (**Extended Data. Fig. 6a**), applying ChAI-Net to the sequences before and after the insertion, and quantifying the difference using 1 - Pearson’s correlation (1-*r;* **Fig. 3a, Methods**). For *de novo* motifs discovered by both models that matched the same TF motif, we used the consensus sequence discovered by ChAI-Net.

Across regulatory region types, we observed that TF motifs captured by both models led to the largest changes in predicted co-accessibility, and motifs only learned by the average accessibility model produced the smallest changes (**Fig. 3b**). Insertions of CTCF, NFY, ZNF143, and SOX2 (with or without POU5F1 or POU2F1) were some of the strongest drivers of co-accessibility, consistent with their high importance in all three types of regulatory elements and in both models. We hypothesize that CTCF, NFY, and SOX2 motifs led to larger predicted changes due to the individual abilities of these TFs to access, modify, and stabilize wider DNA accessibility regions via pioneering^42^ or binding DNA with high affinity and long resident time^43^. We also observed large changes in co-accessibility patterns upon insertion of NRF1 or BANP motifs, which were learned from expressed TSSs.

Visualizing how each inserted motif was predicted to change co-accessibility showed that ChAI-Net has learned that most motifs create a footprint characterized by a protected region (centered blue cross in **Fig. 3c**) flanked on both sides by an average of ∼160 bp of nucleosome-free DNA. In each case, additional subsequences of each window are predicted to increase in their co-accessibility with this central TF-bound region (and with each other). While CWMs from two or three types of regulatory regions often matched the same TF, their consensus sequences were not identical. In several cases, this produced different predicted effects on co-accessibility upon insertion. For example, the enhancer-based NFY sequence was able to alter co-accessibility effectively with flanking nucleosome linker regions, whereas the CTCF-based sequence shows less coordinated accessibility in adjacent linker regions (**Figs. 3b, c**). NFY’s effect was correlated with the GC-content of the background sequences (**Extended Data. Fig. 6b**). For ZNF143, the TSS-based motif shows greater ability to increase co-accessibility at flanking linker regions than does the enhancer-based motif (**Fig. 3c**). Altogether, these results indicate that somewhat different TFs drive co-accessibility as opposed to simply accessibility and that their relative importance varies across different types of regulatory elements.

### GC-rich TF motifs combinatorially regulate enhancer co-accessibility

Having discovered many co-accessibility motifs in enhancers, we were curious what ChAI-Net had learned regarding combinatorial TF binding. For each pair of *de novo* CWMs learned by ChAI-Net, we computed the co-occurrence enrichment across enhancers, with co-occurrence meaning that both motifs are important to predicting co-accessibility in a given enhancer. Then we repeated this for the average accessibility model in order to distinguish which pairs are uniquely important to co-accessibility.

For both models, we observed groups of mutually co-occurring and co-excluding motif pairs (**Fig 4a**). One cluster for each model (average accessibility cluster I, ChAI-Net cluster i) contained many *de novo* motifs with negative contribution scores. Many of these did not match any known TFs, though a few matched NFY, TGIF2, and E2F4 (**Fig. 4b, Extended Data Fig. 3a**). This suggests that enhancers contain pairs of sequences that prevent each other from being accessible on the same chromatin fiber (i.e., one is accessible or the other but not both), as would be expected from proteins competing to bind partially overlapping sites or positioning nucleosomes over flanking regions.

**Figure 4.**
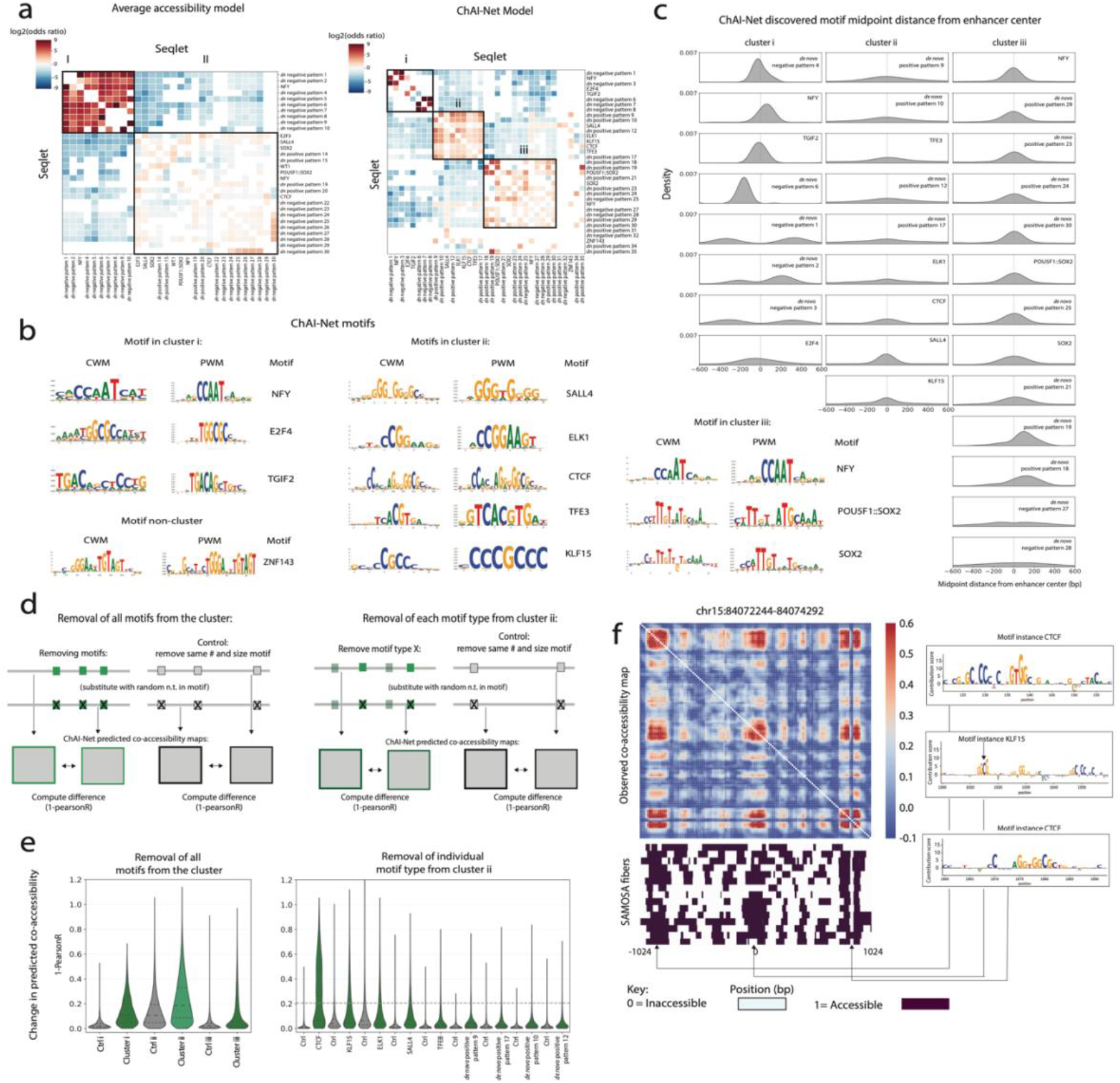
ChAI-Net discovers CpG-rich enhancer motifs. **a**. Co-occurring *de novo* motifs discovered in enhancers by the average accessibility model (left) and ChAI-Net (right), with CWM-PWM matching TFs labeled. Significantly (Fisher’s exact test BH-adjusted *p* < 0.05) enriched (red) or depleted (blue) motif pairs are clustered (I, II, i, ii, iii) based on their co-occurrence patterns across enhancers. **b**. Select enhancer motifs discovered by ChAI-Net: CWM (left), most similar PWM (middle), motif name (right). **c**. Distribution of *de novo* motif instance midpoints relative to the enhancer center (zero). **d**. Schematic of *in silico* motif ablation experiments. Left: replacement of all ChAI-Net motif instances from a given cluster (green boxes) or matched control sequences (grey boxes) with random sequences. Right: ablation of instances of one motif at a time (dark green boxes). **e**. Change in predicted co-accessibility upon motif ablation. Left: Ablating all motif instances from each cluster (shades of green) altered co-accessibility more than ablating matched controls (grey) (MWU test BH-adjusted *p* < 0.05 for all three clusters). Clusters varied in the number of enhancer windows: cluster i (n=1,570), cluster ii (n=22,823), cluster iii (n=16,335). Right: Ablating instances of individual motifs (green) altered co-accessibility more than their matched controls (all MWU test BH-adjusted *p* < 0.05) but generally less than the combination of all motifs (median cluster ii: dashed line). **f**. Example of an enhancer window with co-accessible regions that were sensitive to ablation of the underlying ChAI-Net cluster ii motifs.

Both models had a second group of co-occurring motifs (average accessibility cluster II, ChAI-Net cluster iii) with positive contribution scores. These included many *de novo* motifs without similarity to known TFs, as well as matches to pluripotency TFs like POU5F1::SOX2 and SOX2. NFY also matched *de novo* motifs in these clusters; this TF having both positive and negative CWMs is consistent with previous characterization of NFY bifunctional activation and repression activity associated with transcription and epigenetic marks^44^.

Beyond these shared patterns, the co-accessibility model uniquely identified a subset of enhancers containing co-occurring *de novo* motifs (cluster ii) that were not captured by the average accessibility model and rarely co-occurred with motifs from clusters i or iii. This cluster does not contain matches to POU or SOX family TFs, consistent with a subset of mESC enhancers being distinctly regulated by non-pluripotency TFs ^45^. Some cluster ii motifs matched known TFs, including SALL4 and a subcluster with matches to CTCF, ELK1 (ETS family), KLF15 (SP1 family), and TFE3. Other *de novo* motifs did not match TFs, and several of these exhibited high CpG dinucleotide content. Two GC-rich CWMs were characterized by gaps between invariant bases (*de novo* pattern 9 and 10) that resemble the complex CpG island (CGI)-associated TF motifs found in Codebook^46^ (**Extended Data Fig. 3a**). Since enhancers containing cluster ii motifs resemble distally located (“orphan”) CGIs that exhibit dynamic DNA methylation, increased contact frequencies relative to classical enhancers, and enrichment for CTCF plus SP1 and ETS family motifs^47^, we tested them for overlap with annotated CGIs, observing significant enrichment (Fisher’s exact test, P < 0.001) (**Extended Data Fig. 7b**). Further supporting orphan CGI function, CTCF and TFs with CXXC-domains that recognize CG-rich sequences are sensitive to CpG methylation^46^, and SALL4 and ELK1 are capable of recruiting DNA methyltransferases that modify cytosines^48,49^. Together, these findings highlight a subset of mESC enhancers that harbor motifs for proteins that respond to and modulate local DNA methylation, providing a potential mechanism for the importance of the TFs in co-accessibility.

To better understand the enhancer subtypes identified by co-accessibility analysis, we examined ChAI-Net motif densities and distributions within individual enhancers (**Fig. 4c**). Cluster i (negative CWM) motifs preferentially occurred away from the enhancer center, suggesting a role in positioning nucleosomes and preventing co-accessibility between the centrally bound sequence and flanking DNA. In contrast, cluster iii (positive CWM) motifs showed positional preference towards the enhancer center, indicating their importance in establishing large open chromatin regions at pluripotency TF binding sites. Compared with clusters i and iii, cluster ii motifs occurred in more enhancers and at higher density, with an average of more than one instance per enhancer window (**Extended Data Fig. 7a**) and a broad distribution throughout the enhancer. This is consistent with the higher TF-binding site density reported for enhancer CGIs (∼2–4 sites per kb)^47^. Together, these results support different functional roles for TFs in each ChAI-Net motif cluster.

To further characterize the functions of ChAI-Net motifs, we again leveraged *in silico* experiments. We computationally replaced all instances of each TF motif in enhancer sequences with random non-motif nucleotides from the same enhancer, made ChAI-Net predictions before and after motif ablation, and quantified changes in co-accessibility (**Fig. 4d**). As a negative control for each motif, we repeated this process for equal numbers of length-matched non-motif DNA sequences. Motifs from all three clusters showed more change in co-accessibility upon ablation compared to random non-motif control sequences (**Fig. 4e**), confirming their importance to the sequence patterns learned by ChAI-Net.

To understand the joint importance of co-occurring motifs, we next conducted an experiment ablating all instances of all motifs from cluster i, ii, or iii. Cluster ii motifs showed the largest effect on co-accessibility (**Fig. 4e**), which exceeded the average effects of individual cluster ii motifs. However, a small number of enhancers had co-accessibility patterns that were highly sensitive to ablation of all CTCF motif instances, similar to what we observed for CTCF insertions (**Fig. 3b**). Consistent with driving co-accessibility throughout the enhancer window, examples of cluster ii motif ablation within individual enhancers showed large predicted reductions in co-accessible sites and nucleotides with high contribution scores in both central and flanking regions that are jointly accessible on the same fibers (**Fig. 4f**). These findings suggest that cluster ii enhancers encode diverse TF motifs that co-bind and drive coordinated accessible sites over distances of up to 2 kb.

As expected, ablating of all cluster i motif instances had a much larger effect than removing instances of single motifs (**Extended Data Fig. 7c**), suggesting that most *de novo* motifs with negative contribution scores cooperate to repress enhancers. However, a motif matching TGIF2 (from 288 enhancers) and *de novo* negative pattern 3 (from 166 enhancers), discovered only by ChAI-Net, had notable effects on predicted co-accessibility. This suggests that these motifs repress chromatin accessibility across genomic distance, consistent with the existing role of TGIF2 as a repressor of neuronal differentiation by recruiting histone deacetylase corepressor complexes^50^.

In sum, ChAI-Net adds a number of *de novo* enhancer motifs regulating chromatin co-accessibility to our current TF-centric understanding of DNA-protein interactions, highlighting the importance of co-binding in a prevalent set of CGI-like distal enhancers.

### Ruler TFs co-bind to regulate accessibility at active promoters

Across expressed TSSs, ChAI-Net discovered both motifs that are shared with enhancer regions and others that are unique to transcription start sites. Motifs similar to those discovered in enhancers included CG-rich motifs suggestive of CGI-based regulation and matches to CTCF, ZNF143, NFY, ETS family, SP family, NRF1, and POU5F1::SOX2 (**Extended Data Fig. 3b**), which were also detected by modeling average accessibility at active TSSs (**Extended Data Fig. 5b**). Additional TSS *de novo* motifs that were not detected at enhancers included matches to CRE and BANP, with BANP only being detected by ChAI-Net and not the active TSS average accessibility model (**Extended Data Fig. 8**).

The majority of ChAI-Net’s active TSS CWMs matched the motifs of “ruler” TFs (rTFs; **Fig. 5a**) that contribute to determining transcription start locations and regulating transcription levels^51– 58^. Because rTFs share a common feature of glutamine-rich and hydrophobic intrinsically disordered activation domains that form a Velcro-like environment, these TFs flexibly cooperate, physically interact with TFIID, constrain the position of the pre-initiation complex (PIC) during its promoter recognition phase, and contribute to transcriptional activation^51 59–61^. To evaluate if ChAI-Net is jointly leveraging motifs of multiple rTFs to predict co-accessibility at individual TSSs, we examined the prevalence, distribution, and co-occurrence patterns of each rTF’s motif instances. SP, NRF1, and BANP motifs were the most prevalent, with on average >1 instance per TSS (**Fig. 5b**). The rTF motifs showed high co-occurrence with each other, but not with the remaining *de novo* motifs that were also not classified as rTFs: CTCF and POU5F1::SOX2 (**Fig. 5c)**. NFY, CRE, ZNF143, and ETS motifs tended to occur near or upstream of the TSS, while NRF1, SP, and BANP motifs showed broader positioning with a preference downstream of the TSS (**Fig. 5d**). These findings suggest that many expressed TSSs in mESCs are guided by cooperative binding of multiple rTFs over a range of distances up and downstream of the PIC.

**Figure 5.**
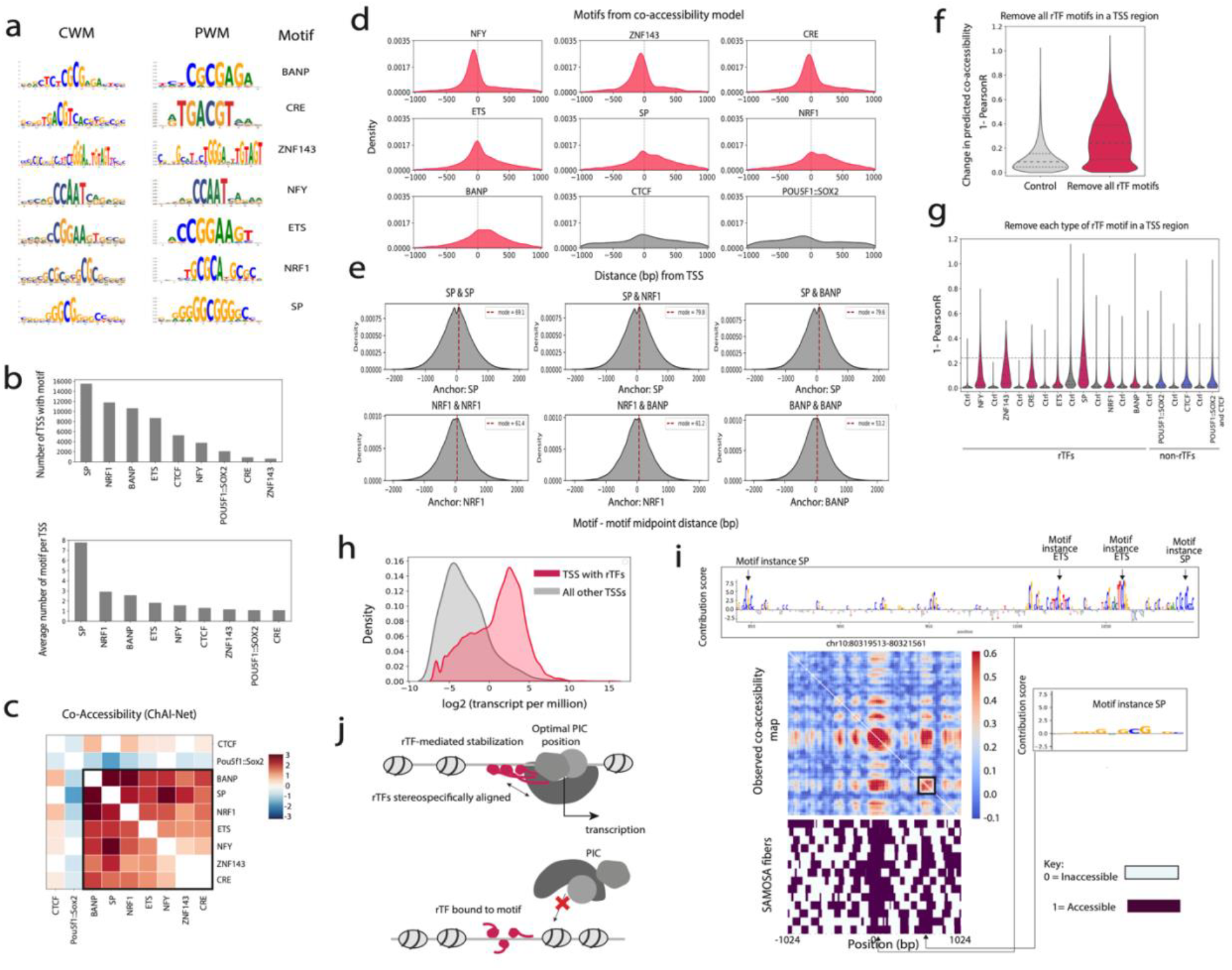
ChAI-Net learns the spacing of ruler transcription factors (rTFs) at transcription initiation sites. **a**. Select *de novo* motifs discovered by ChAI-Net in expressed TSSs: CWM (left), most similar PWM (middle), PWM name (right). **b**. Number of expressed TSSs harboring each ChAI-Net motif (left) and average number of motif instances per 2,048-bp TSS window (right). **c**. Co-occurring *de novo* motifs discovered in expressed TSSs by ChAI-Net, with CWM-PWM matching TFs labeled. Significantly (Fisher’s exact test BH-adjusted *p* < 0.05) enriched (red; black box) or depleted (blue) motif pairs are clustered based on their co-occurrence patterns across TSSs. **d**. Distribution of *de novo* motif instance midpoints (rTFs: magenta, other motifs: grey) relative to the TSS (zero, all oriented 5’ to 3’ left to right). **e**. Motif-to-motif instance midpoint distances for representative pairs of rTFs that co-occur in TSSs. The red dashed line indicates the most common value per pair. **f**. Ablation of all rTFs motif instances (magenta) altered co-accessibility more than ablation of matched control sequences (grey; MWU *p* < 0.05; n=16,437 windows). **g**. Ablation for individual rTFs (magenta) and other TFs (blue) altered co-accessibility more than matched controls (grey; MWU with BH-adjusted *p* < 0.05). Horizontal dashed line: median effect of ablating all rTF motifs. **h**. Expression level per TSS (transcripts per million) stratified by those harboring rTF motifs (magenta) versus not (grey). **i**. Example TSS window with co-accessible sites that are sensitive to ablation of rTF motifs. **j**. Model of rTF stereospecific alignment and Q-rich activation domain cooperativity suggested by ∼10-bp spacing of ChAI-Net CWM instances (top). Such an orientation would mediate stability of the preinitiation complex (PIC) and favor TSS activity, whereas irregular spacing and orientation of rTFs would not (bottom).

To further investigate rTF motif cooperativity, we analyzed the midpoint distances of co-occurring rTF motifs in the same TSS window. We observed that SP, NRF1, and BANP motif instances were consistently spaced ∼60, ∼70 or ∼80 bp apart (**Fig. 5e**). Because DNA rotates approximately 10.5 bp per turn, this motif periodicity suggests these rTFs have flexible spacing but are typically positioned on the same face of DNA, potentially facilitating TF cooperativity and PIC positioning through stereospecific alignment^51^.

Because we obtained these results using CWMs from our co-accessibility model, they are implicitly linked to coordinated binding. To test this more explicitly, we next quantified the importance of TF motifs for co-accessibility at TSSs using *in silico* ablations. For each active TSS with co-occurring rTF motifs, we performed motif ablation by substituting random non-motif nucleotides and predicting the effect on co-accessibility. Simultaneous ablation of all rTF motifs had large effects across expressed TSSs, exceeding the effects of equal numbers of length-matched negative control (non-motif) sequences (**Fig. 5f**). Individual rTFs also had notable effects on co-accessibility, exceeding those of POU5F1::SOX2 and CTCF (**Fig. 5g**), though no single rTF reached the level of change observed when ablating all rTFs per TSS window.

Linking these results to gene expression, TSSs with rTF motifs were expressed at higher levels than other TSS (**Fig. 5h**). Within individual loci that were sensitive to ablation of rTF motifs, we observed strong agreement between co-accessible regions and the locations of high contribution scores that match rTF motifs (**Fig. 5i**). Together, our results support a model in which a mix of different rTFs bind on the same face of DNA to cooperatively regulate transcription from a range of distances using their disordered activation domains (**Fig. 5j**).

### ChAI-Net discovers motifs influencing co-accessibility adjacent to CTCF binding sites

Although CTCF is a major determinant of chromatin organization, the sequence features that distinguish accessible and occupied CTCF sites remain unclear^25^. Given evidence that neighboring DNA-binding proteins can modulate CTCF occupancy and insulation in a cell-type-specific manner^62^, we hypothesized that ChAI-Net would identify the motifs of co-factors that contribute to co-accessibility at CTCF bound regions and that these might differ from the motifs learned at enhancers and TSSs.

Beyond the expected CTCF and pluripotency TF motifs, ChAI-Net discovered *de novo* motifs with positive contribution scores matching TFs known to colocalize with CTCF, including E2F3, NFY, and ZNF143^54,62,63^ (**Fig. 6a, Extended Data Fig. 3c**). E2F3, a regulator of cell cycle and differentiation^63^, was not identified by an average accessibility model (**Extended Data Fig. 5c, Extended Data Fig. 9a**) but was particularly important to ChAI-Net predictions, being present in over 10,000 CTCF bound regions with an average of ∼8 copies per region (**Extended Data Fig. 9b**). ChAI-Net also revealed motifs predicted to decrease co-accessibility at CTCF sites. These included VDR, known to recruit repressive complexes^64^; ZNF384, a CTCF co-regulator associated with repressing DNA accessibility flanking CTCF bound sites ^39^; and *de novo* motifs 4 and 5 that were highly prevalent but did not match a known TF (though *de novo* motif 4 is somewhat similar to the YY1 motif). Most of these motifs were not discovered by a CTCF site average accessibility model, which primarily identified CTCF and pluripotency TF motifs (**Extended Data Fig. 5c**).

Evaluating the co-occurrence of *de novo* motifs within CTCF sites revealed two distinct clusters co-occurring motifs (**Fig. 6b**), which were not prominently observed from the average accessibility model (**Extended Data Fig. 9a**). In cluster i, motifs matching SOX2, E2F3, and *de novo* negative motifs co-occurred, with only E2F3 showing positional preference for binding near CTCF and at periodic preferred distances (**Fig. 6c, Extended Data Fig. 9c**). This suggests that E2F3 may bind in the linker DNA of phased nucleosomes flanking CTCF or possibly at the edges of nucleosomal DNA, which has been observed *in vitro* for E2F1^65^. On the other hand, cluster ii was characterized by co-occurrence of NFY, ZNF384, and/or VDR motifs. ZNF384 and VDR were enriched flanking CTCF, while NFY tended to bind just upstream of CTCF, possibly indicating a physical interaction. Supporting the idea that ChAI-Net identified CTCF co-factors, CTCF insulation can depend on sequences flanking its binding site^66^, and NFY can promote local chromatin accessibility and exhibits constrained positioning with neighboring TFs^67^. We did not observe different orientations or lengths of CTCF motifs to co-occur with each other at the ∼2-kb window scale. Instead, our results highlight that CTCF bound sites are co-accessible with adjacent sites bound by one of several different sets of co-occurring TF combinations at preferred distances.

**Figure 6.**
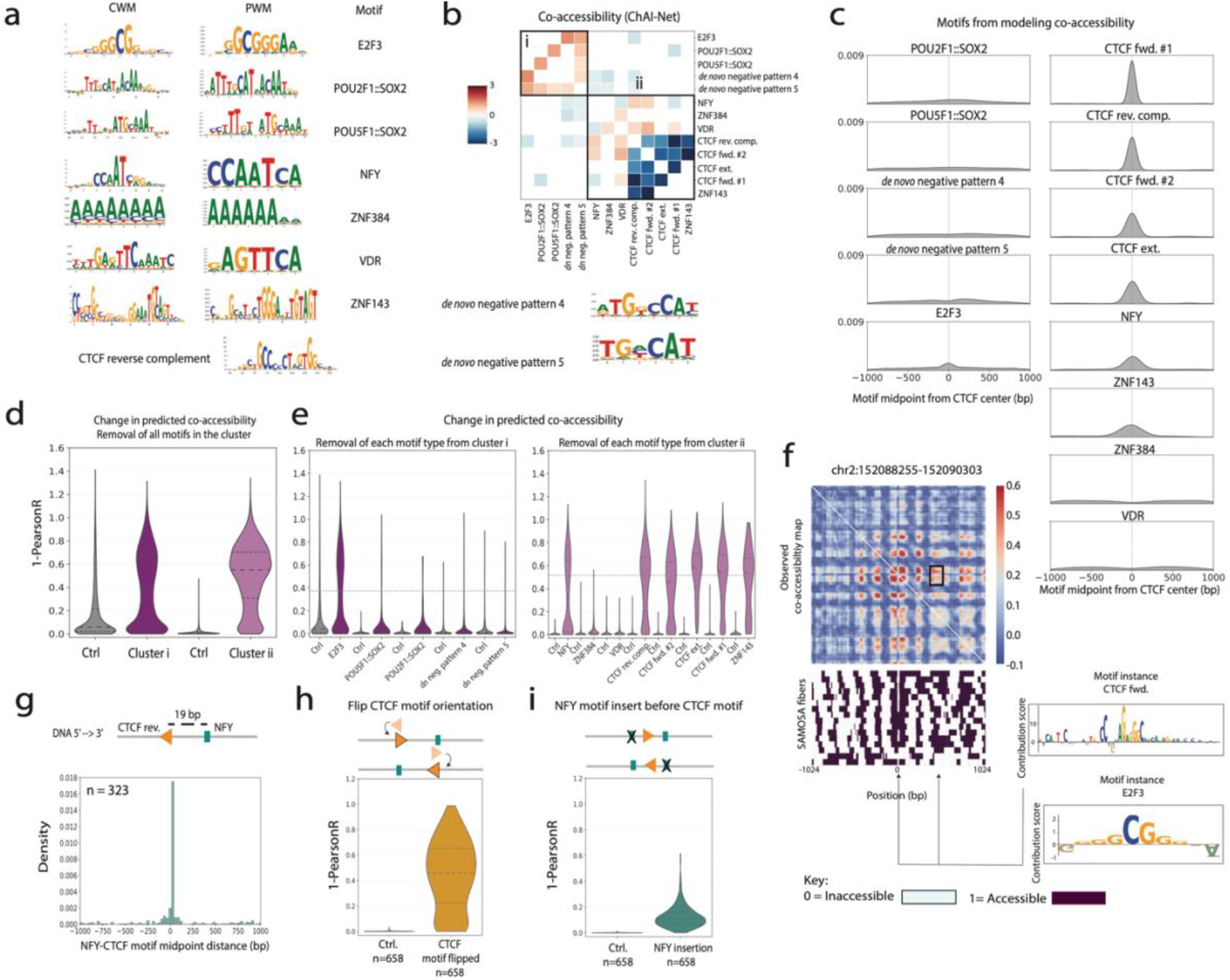
ChAI-Net learns that E2F3 and NFY differentially influence co-accessibility at CTCF sites. **a**. Select motifs discovered by ChAI-Net in CTCF sites: CWM (left), most similar PWM (middle), PWM name (right); CWM only for CTCF and *de novo* motifs without PWM matches. Additional motifs in Extended Data. Fig. 3C. **b**. Co-occurring *de novo* motifs, with CWM-PWM matching TFs labeled. Significantly (Fisher’s exact test BH-adjusted *p* < 0.05) enriched (red) or depleted (blue) motif pairs are clustered (i, ii) based on their co-occurrence patterns across CTCF sites. **c**. Distribution of *de novo* motif instance midpoints relative to the CTCF peak center (zero). Left column: cluster i motifs; Right column: cluster ii motifs. **d**. Ablation of all motif instances per cluster (shades of purple) altered co-accessibility more than ablation of matched control sequences (grey; MWU *p* < 0.05; n=5849 for cluster i windows, n=11014 cluster ii) **e**. Ablation for individual motifs from cluster i (dark purple) or cluster ii (light purple) altered co-accessibility more than matched controls (grey; all MWU BH-adjusted *p* < 0.05). Horizontal dashed line: median effect of ablating all motifs in the cluster. **f**. Example CTCF site with co-accessible sites that are sensitive to ablation of cluster i motifs. **g**. Motif-to-motif midpoint distances for CTCF reverse complement and NFY instances co-occurring in the same window (n=323). The most common distance is labeled in the cartoon above the histogram. **h**. Flipping the orientation of CTCF forward or reverse complement motifs altered co-accessibility significantly more than flipping matched control sequences (MWU BH-adjusted *p* < 0.05; n: total windows) **i**. Ablating NFY motif instances and inserting a NFY consensus sequence at a motif-to-motif midpoint distance of 19 bp in front of the CTCF forward or reverse complement matching motif altered co-accessibility significantly more than performing these experiments with a matched control sequence (MWU BH-adjusted *p* < 0.05; n: total windows).

To test the contribution of each discovered motif to predicted co-accessibility, we performed *in silico* motif instance ablation (**Methods**). Motifs in cluster i and cluster ii showed similar magnitude effects when ablated simultaneously, and both groups were significantly greater than their respective negative controls (**Fig. 6d**). Ablation of E2F3 (cluster i) and NFY (cluster ii) motifs showed effects nearly the same as ablating all motifs in their cluster (**Fig. 6e**), as did ablation of various CTCF motifs. Ablation of all E2F3 motifs had a much large effect than ablation of individual motif instances (**Fig. 3b**), indicating cumulative effects for this highly abundant motif. Consistent with this, CTCF sites that were most sensitive to E2F3 motif ablation had very high co-accessibility showing patterns of periodic jointly open chromatin regions (**Extended Data Fig. 9d**), which often corresponded to the E2F3 motif locations (**Fig. 6f**). Other individual motifs had smaller effects, though most exceeded their negative controls. These results demonstrate that E2F3 is particularly important in determining co-accessibility at CTCF sites.

Another important ChAI-Net motif corresponded to NFY, which co-occurred with ∼6% (n=335) of CTCF forward-matching motifs and ∼10% (n=323) of CTCF reverse-matching motifs (**Extended Data Fig. 9e**). NFY instances had a preferred position ∼19 bp away from the CTCF binding site (**Fig. 6c**) and their orientation depended on the orientation of CTCF (**Fig. 6g, Extended Data Fig. 9f**). To explore if these relationships are functional, we flipped the orientation of the CTCF motifs next to NFY instances, observing large predicted effects on co-accessibility (**Fig. 6h, Methods**) that removed a pattern corresponding to phased nucleosomes flanking the CTCF binding site (**Extended Data Fig. 9g**). We observed additional changes in predicted co-accessibility at these loci upon replacing native NFY motif instances with the ChAI-Net NFY consensus sequence (**Fig. 6i, Methods**), including a slightly wider and stronger co-accessible region flanking the CTCF motif (**Extended Data Fig. 9g**). Finally, we inserted extra NFY consensus sequences into these loci at distances >20 bp from CTCF, observing large effects only when insertions aligned with open, co-accessible regions (**Extended Data Fig. 9h, Methods**). These experiments suggest that the phased nucleosomes typically flanking CTCF sites constrain the binding of NFY to accessible linker DNA or possibly that NFY plays a role in positioning nucleosomes (**Extended Data Fig. 9i**).

Together, these results pinpoint E2F3 and NFY as TFs that colocalize with CTCF, interacting with CTCF in two distinct ways to promote co-accessibility.

## Discussion

Here, we introduce ChAI-Net, a deep learning model designed to learn sequence elements contributing to simultaneous accessible (co-accessible) regions across kilobase pairs of DNA. Unlike existing models^20–22,68–71^ designed to predict 1D epigenomic signals, ChAI-Net combines an Akita-like framework^33^ with the unique information captured by assays like SAMOSA to predict 2D chromatin co-accessibility. Trained on more than 10 million genomic regions, ChAI-Net enabled us to extract *de novo* sequence motifs that are critical for accurately predicting co-accessibility in enhancers, expressed TSS, and CTCF sites. It also provides a framework to then perform *in silico* experiments to establish if observed relationships amongst these motifs causally affect predicted co-accessibility patterns. These analyses suggested that co-accessibility is guided by a set of *de novo* motifs that vary across regulatory contexts, many of which are not discoverable through modeling an average representation of chromatin accessibility.

Key insights afforded by ChAI-Net include specific motif pairings, orientations, and spacings underlying coordinated TF occupancy in regulatory elements^3^. Within a subset of enhancers, we identified CpG-rich motifs consistent with DNA methylation based regulation involving Polycomb, Mediator and/or ZF-CXXC proteins^46^, including motifs matching TFs known to be sensitive to DNA methylation^46^, enriched in CGIs, or capable of recruiting methylation modification machinery^48,49^. While not observable in ∼2-kb windows, these TFs may also contribute to the selective enhancer– promoter communication via homotypic CGI-CGI interactions^72^. Among expressed TSSs, we discovered motifs matching rTFs^51^, and our analyses confirmed the importance of rTF co-binding suggestive of stereospecific alignment on the same face of DNA in order to utilize the long activation domains to optimally position the PIC^51^. At CTCF sites, ChAI-Net highlighted periodic E2F3 co-binding between nucleosomes and NFY binding at ∼19 bp from CTCF in a conserved orientation that is consistent with previous ChIP-seq analyses^73^. These relationships amongst TFs are obscured in bulk data but were learnable with ChAI-Net specifically because we chose to model co-accessibility. Together, these data identify a subset of TFs for which binding to regulatory elements is not an independent event, but rather one that is coordinated with other proteins along individual chromatin fibers and consistent with cooperative relationships in carrying out important nuclear processes, as previous described^3,74^.

These results underscore that ChAI-Net and SAMOSA data can facilitate discoveries beyond what is observable without deep learning or from bulk measurements. PWM analysis is challenged by the fact that many potential binding sites are not occupied. ChIP-seq, ATAC-seq and other bulk genomic assays help to narrow sequence motif instances to those more likely to be regulatory, but they are limited in resolution. For example, peak intersections show that two proteins bind nearby but cannot distinguish if they bind simultaneously versus exclusively or randomly, and multiple nearby binding events can be difficult to disentangle. Single-molecule footprinting with SAMOSA or related techniques addresses these gaps by measuring protein-DNA interactions on individual chromatin fibers at near-bp resolution. Deep learning with a model such as ChAI-Net provides another layer of information, because it quantifies how important individual nucleotides are to the prediction task. In this case, by predicting co-accessibility, this strategy reveals which sequences are most important to reconstructing co-binding events. Matching these *de novo* co-accessibility motifs to TF PWMs clarifies which DNA-binding proteins co-bind, while the remaining motifs not matching a known TF are useful for generating hypotheses about new mechanisms, such as DNA methylation-based regulation.

Despite underscoring the importance of cis-regulatory codes in determining chromatin architecture, our study has several limitations. A first set of caveats relates to the SAMOSA data used to train ChAI-Net. Compared to bulk approaches (e.g., ATAC-seq, DNase-seq, ChIP-seq), all genome-wide single-molecule footprinting methods are limited by read coverage^9^, making it challenging to detect events that are highly dynamic or transient. In this study, low coverage led to some loci being filtered out, and it limited our modeling to ∼2-kb windows, which ensured sufficient window-spanning reads for robust co-accessibility calculations at near-bp resolution. This means that long-range co-accessibility, such as between distal enhancers and their target promoters, could not be investigated. New technologies that combine footprinting with chromatin capture and/or longer reads promise to resolve this limitation (e.g., SMF-ONT, LASSI^74,75^), and it will be exciting to extend the ChAI-Net approach to these emerging data.

Second, we focused on mESCs due to high SAMOSA data quality and coverage, as well as previously validated annotations of regulatory elements. However, further training on co-accessibility measurements from a broad range of cellular contexts would enable a deeper understanding of the sequence grammar underlying chromatin co-accessibility, including elucidation of new combinations of regulatory motifs and insight into which of our observations are conserved across species, cell types, and developmental stages.

Lastly, ChAI-Net is a powerful tool for extracting regulatory sequence rules and computationally testing causal hypotheses regarding the sequence elements guiding co-accessibility. But these predictions require experimental validation with methods such as genome and epigenome editing combined with targeted single-molecule footprinting. These technologies are not yet robust enough to test the multitude of hypotheses generated by ChAI-Net at scale. But we look forward to future work exploring the coupling of deep learning with laboratory testing. Meanwhile, ChAI-Net is a promising strategy for generating and testing mechanistic hypotheses *in silico* and prioritizing TFs and binding sites for downstream experiments.

As single-molecule footprinting approaches continue to advance, ChAI-Net represents an example of how a deep learning model can be applied to these datasets to reveal and disentangle TF binding partners.

## Methods

### Co-accessibility data

SAMOSA reads from mESCs (GSE245558) were processed using a neural network hidden Markov model, as previously described^23^. On each sequenced molecule, nucleotide level accessibility calls above 0.5 were set to 1 (accessible nucleotide) and below 0.5 to 0 (not accessible nucleotide).

The mouse genome (mm10 assembly) was divided into nonoverlapping virtual contigs and randomly assigned to training, validation, and test sets with an 80/10/10 split, as previously described^33^. We extracted 2,048-bp sequences using sliding genomic windows with 75% overlap for the training set and 50% overlap for the validation and test sets. This procedure resulted in ∼4 million training, ∼243 thousand validation, and ∼243 thousand test examples. These sequences were the “features” (inputs) for ChAI-Net training.

The coordinates of these DNA sequences (“windows”) were used to extract corresponding SAMOSA fibers aligned to mm10. We retained windows that had at least 8 SAMOSA fibers that spanned the entire 2,048 bp window, because contiguous coverage is important for quantifying co-accessible sites. Co-accessibility was calculated for each pair of nucleotides by counting the proportion of aligned fibers on which both nucleotides are accessible. For training ChAI-Net, we averaged these values into 16-bp x 16-bp bins, creating a symmetric matrix of 128 × 128 bins. Then, we normalized each of these matrices by subtracting an estimate of the expected matrix in the absence of open chromatin regions and well positioned nucleosomes. This estimate was calculated using the bin-wise mean of matrices for all windows in the fold (i.e., separately for training, validation, and test to prevent leaky pre-processing). The resulting normalized (observed minus expected) co-accessibility maps were the “targets” (outputs) for ChAI-Net modeling.

To generate a corresponding average accessibility track for each window, we computed the mean accessibility across fibers per nucleotide in the window and then averaged these values into 16-bp bins, generating a vector of length 128 bins per window. No normalization was performed.

### Estimation of genome-wide sequencing coverage for SAMOSA data

Genome-wide sequencing coverage was quantified from aligned BAM files using mosdepth (v0.3.3). For each BAM file, mean sequencing depth was calculated across the mouse mm10 reference genome, and coverage breadth was defined as the fraction of genomic bases covered by at least one (≥1x) or two (≥2x) aligned reads. Coverage metrics were calculated independently for each of the BAM files and subsequently summarized across files.

### Visualization of publicly available epigenomic data

For visualization, publicly available mESC H3K27ac ChIP–seq and ATAC-seq signal tracks were obtained from the 4D Nucleome (4DN) Data Portal^76^. H3K27ac data were obtained from experiment set 4DNESDVQP5XT (processed signal fold-change bigWig file 4DNFIHRTWNB4), and ATAC-seq data were obtained from experiment set 4DNESQIDKLL7 (processed file 4DNFIIY4QW2D; GEO accession GSE198517)^77^. These data were used for visualization only and were not used for model training or quantitative analyses.

### Model architecture for ChAI-Net and average accessibility models

We developed the ChAI-Net architecture and training strategy by modifying Akita, a convolutional neural network that predicts two-dimensional (2D) Hi-C maps from one hot encoded DNA sequences^33^, to predict co-accessibility. ChAI-Net consists of a ‘trunk’ that has a similar architecture as Akita but modified to accept 2,048-bp input sequences and then utilize a smaller 1D convolution tower to arrive at a vector of length 128 representation of the input (corresponding to the 128 × 128 binned co-accessibility targets). The trunk’s role is to estimate 1D representations of input DNA sequences (features). This is performed by scanning each window’s one-hot encoded sequence (presented to the model in batches) to learn DNA patterns and their combinations, which are compressed into a representation of the input sequence that is the output of the trunk and the input to a ‘head’ that predicts co-accessibility. The head of ChAI-Net transforms the 1D representation into a 2D co-accessibility map by learning relationships between compressed DNA features, combined across genomic distance through multiple dilated residual 2D convolutions, and the values in the co-accessibility maps (targets). Training estimates 421,617 trainable parameters across the trunk and head that optimize the mean squared error (MSE) between the upper triangle of predicted targets versus the upper triangle of the true co-accessibility targets.

To predict average accessibility, we retained the same ‘trunk’ architecture and paired it with a width-one convolutional layer to predict the 1D average accessibility tracks (targets). Wee used the same training/validation/test sets. The average accessibility model has 323,905 total trainable parameters, which are estimated by optimizing MSE between predicted and true average accessibility targets.

We implemented both models using the Basenji software, which was written in TensorFlow and Keras (https://github.com/calico/basenji/tree/master/manuscripts/akita).

### Training approach

For ChAI-Net, we fit the model parameters using stochastic gradient descent with momentum for ∼60 epochs. Data augmentation was used to avoid overfitting and maximize generalization accuracy to unseen sequences. For each input DNA sequence, we stochastically shifted input sequences by up to +/-11 bp and reverse complemented the DNA and flipped the co-accessibility map or average accessibility track. Following the Akita framework, we stopped training when validation loss did not improve for 12 epochs, and retained the model the epoch with minimum MSE loss on the validation set. We tried several learning rates (0.0001, 0.0065, 0.001, 0.01) and batch sizes (2, 4, 16 windows) to identify values with the best performance, which was achieved with a learning rate of 0.0065 and batch size of 16 windows. For average accessibility, we used the same hyperparameters, data augmentation, and optimization strategy.

### Regulatory regions used for model interpretation

TSSs were based on their gene-level expression estimates from mouse ES-E14 cells obtained from the ENCODE portal (accession ENCSR000CWC; file ENCFF827OZU)^26–28^. Expressed TSSs were defined by taking TSSs of genes with a TPM greater than 0, while non-expressed TSSs were defined as genes with a TPM equal to 0. We used previously identified enhancers classified as promoter-distal ATAC-seq peaks with associated PRO-seq signal^29^. We used previously identified ChIP-seq defined CTCF binding sites from ENCODE with high Find Individual Motif Occurrences (FIMO) scores^30,31^. For each regulatory element, we generated the DNA sequence and normalized co-accessibility map of a 2,048-bp window centered on the element, using peak centers for enhancers and CTCF sites. We also made predictions and computed importance scores (described below) for each regulatory element centered genomic window.

### Model performance evaluation on held-out test data overlapping regulatory regions

To assess performance of ChAI-Net and the average accessibility model, we made predictions for all windows in the held-out test set. Additionally, we made predictions for all 2,048-bp windows centered on an enhancer, expressed TSS, non-expressed TSS or CTCF site with at least 97% overlap with a held-out test set window, computed using BEDTools (v2.31.1) intersect (-wa -f 0.97). The remaining held-out test windows were labeled as non-regulatory if they had less than 97% reciprocal overlap with any of the regulatory element centered windows (-v -r -f 0.97). For each of the resulting regulatory or non-regulatory windows, we used MSE and Pearson’s correlation (*r*) to compare the observed and predicted co-accessibility map. These statistics were then used to evaluate the performance of ChAI-Net. Similarly, we used MSE and *r* to compare observed and predicted average accessibility tracks for each window to evaluate the performance of the average accessibility model.

### Quantification of observed co-accessibility score

To quantify the degree of co-accessibility in each observed co-accessibility map, values within two bins of the main diagonal were first set to zero to exclude near-diagonal interactions. An 8 × 8 sliding filter was then applied across the co-accessibility map, and the mean co-accessibility within each filtered region was calculated. The observed co-accessibility score was defined as the maximum of the resulting values. This score captures the largest off-diagonal co-accessible region per window, representing a distal set of bins that are most frequently open on the same fibers.

### ChAI-Net and average accessibility model interpretation

We performed model interpretation to determine the extent to which each nucleotide was predictive of chromatin co-accessibility or average accessibility. We interpreted regulatory regions (i.e., windows centered on expressed TSSs, enhancers, and CTCF sites). We used the DeepLIFT algorithm^37^ to compute contribution scores for each nucleotide in the 2,048 bp input windows with respect to the predicted co-accessibility map or average accessibility track. Contribution scores were derived for each example from each regulatory region using either ChAI-Net or the average accessibility model.

### Discovery of *de novo* motifs and identification of predictive motif instances

Using the contribution scores from either ChAI-net or the average accessibility model, we applied TF-MoDISco-lite^38^ to identify seqlets: short spans of contiguous high positive-importance or high negative-importance nucleotides. This gave rise to six distinct sets of seqlets for expressed TSSs, enhancers, and CTCF sites, from both ChAI-Net and the average accessibility model. For each set, seqlets were clustered, aligned, and used to build *de novo* motif contribution weight matrices (CWMs). Parameters used for running TF-MoDISco included a window size (-w) of 2,048 around the center of each input region and 20,000 as the maximum of positive/negative seqlets (-n).

For annotation, we used TOMTOM^78^ to compute similarities between the *de novo* motif CWMs and a curated set of known transcription factor (TF) binding site position weight matrices (PWMs) from JASPAR^34^ and HOCOMOCO ^35^. For each motif, we manually inspected the most similar PWMs between the databases and assigned the name of the TF with the best PWM match.

To identify genomic instances of de novo motifs, we used the Fi-NeMo package (https://github.com/kundajelab/finemo_gpu, v0.23, commit b81876d), which fits a sparse linear regression model per genomic region to minimize the difference between the contribution scores and a reconstruction of the scores from a weighted combination of trimmed CWMs. When identifying hits in input regions using TF-MoDISco CWMs, we used default parameters for the L1 regularization weight determining the sparsity of hits and for batch size. Rather than using the default universal threshold to determine motif start and end positions, we specified the start and stop positions of each CWM by manually inspecting each *de novo* CWM motif.

### *In silico* insertion of motifs into GC-controlled background DNA sequences

To assess the effects of *de novo* motifs on predicted chromatin co-accessibility, we performed *in silico* motif insertion experiments separately for motifs identified in each regulatory context (enhancers, expressed TSSs, CTCF sites). For *de novo* motifs in each regulatory context, 100 background sequences were generated by sampling 2,048-bp genomic windows and then scrambling them using BiasAway (v3.3.0)^79^, which performs k-mer shuffling within a sliding window. We shuffled nucleotides within a 100 bp sliding window (50 bp step) while preserving the local dinucleotide (k=2) composition. We used the default parameters and a fixed random seed (seed=8) for reproducibility. The consensus sequence of the *de novo* motif’s CWM was subsequently inserted into each of the 100 scrambled background sequences, and sequences with versus without the insertion were input into ChAI-Net to predict co-accessibility. The resulting two predicted maps were used to assess the effect of the motif insertion as the motif effect on co-accessibility (1-Pearson *r* between the two predicted maps). Each inserted *de novo* motif gave a hundred scores (1-*r*) that measured the change in predicted co-accessibility as a result of the motif insertion. We additionally visualized the predicted co-accessibility differences as heatmaps for each motif by subtracting the predictions of the inserted motif against the predictions without the insertions. The same procedure was repeated using the average accessibility model.

### Correlation of GC content with *de novo* motif insertion scores

For each background DNA sequence used for *de novo* motif insertion, GC content was calculated as the proportion of guanine (G) and cytosine (C) nucleotides relative to the total sequence length. The association between GC content and the change in predicted co-accessibility as a result of inserting the *de novo* NFY matching motif (discovered from either enhancer or expressed TSSs sequences) was assessed using both Pearson’s correlation coefficient (*r*) and Spearman’s rank correlation coefficient (*ρ*). Statistical significance was evaluated using two-sided tests, with *P* < 0.05 considered statistically significant.

### Motif co-occurrence analysis

Among *de novo* motifs discovered by ChAI-Net and the average accessibility model, we tested whether instances of two *de novo* motifs co-occurred within regulatory elements more frequently than expected by chance. We assessed each regulatory element-centered window for whether it had each *de novo* motif present or not. Then for each pair of motifs, we populated a 2 × 2 contingency table with the number of windows containing both motifs, one motif, or neither motif. We performed a Fisher’s exact test to calculate a P value for enrichment. We corrected for multiple hypothesis testing across all pairs, using the Benjamini-Hochberg (BH) method. Significant co-occurring motif pairs were defined as those with BH adjusted P values below 0.05.

### *In silico* removal of motif instances

To assess the effect of individual motif instances, we performed *in silico* motif removal by replacing each motif instance with nucleotides randomly sampled from positions within the same window that did not overlap a motif instance. As a control, we selected positions within the 2,048-bp window that did not overlap the motif instances used for removal and that matched the targeted motifs in both number and length (grey boxes with black outlines, **Fig. 4d**). These control positions were similarly replaced with randomly sampled nucleotides from the DNA sequence (grey boxes with black crosses, **Fig. 4d**). ChAI-Net was used to predict a co-accessibility map for each perturbed sequence, and the effect of each perturbation was quantified by comparing the predicted map with that of the corresponding unperturbed sequence using 1 − Pearson’s *r*.

### Enrichment analysis of enhancers and CpG islands (CGIs)

To test for the enrichment of enhancers with ChAI-Net cluster ii motifs overlapping CGIs, we performed a Fisher’s exact test using a 2×2 contingency table constructed from interval-level overlaps. CGI coordinates were obtained from the genomic location of CXXC Affinity Purification (CAP) identified CGIs^80^ and the UCSC Genome Browser CpG Islands track from the mm10 genome build^81^. Overlaps between enhancers with cluster ii motifs and CGIs were determined using BEDTools (v2.31.1) intersect, requiring ≥1 bp of overlap. The background set consisted of enhancers with cluster i and cluster iii motifs without cluster ii motifs, from which the number of CGI-overlapping and non-overlapping intervals was calculated in the same fashion as cluster ii motif containing windows. Statistical significance was assessed using a two-sided Fisher’s exact test implemented in Python (v3.9.16) using SciPy (v1.11.4; scipy.stats.fisher_exact). Odds ratios and 95% confidence intervals were reported. A P value < 0.05 was considered statistically significant.

### *In silico* perturbations for regions centering the CTCF bound site

In CTCF sites with at least one NFY motif instance, we performed several *in silico* sequence perturbation experiments. First, the CTCF motif instance was flipped to be oriented in the opposite direction, keeping the rest of the DNA sequence unmodified. Second, the endogenous NFY motif was ablated by replacing it with non-motif instance overlapping DNA sequence from the same window. The consensus sequence of the NFY CWM identified across CTCF sites by ChAI-Net was then inserted at specified distances upstream of the CTCF motif instance. For Fig. 6I, the NFY motif was inserted in front of the CTCF motif instance with a motif-to-motif midpoint distance of 19 bp. For Extended Data Fig. 9H, the NFY motif was inserted at positions ranging from 25 to 1,000 bp upstream of the CTCF motif in 5-bp increments. For each experiment, ChAI-Net was used to predict co-accessibility maps for the window before and after the *in silico* perturbation. The two maps were then compared using 1 − Pearson’s *r* to quantify the effect of the perturbation.

## Data Availability

All data used in this study is publicly available.

## Code Availability

ChAI-Net and the average accessibility model used to model SAMOSA data are located at https://github.com/jlee-86/ChAI-Net. Code for individual ISM experiments is forthcoming and will be available at https://github.com/jlee-86/ChAI-Net.

## Author Contributions

J.L., V.R., and K.S.P. conceived this study. J.L. created and trained models with the help of K.Z. and E.S., performed computational analyses, and wrote the initial draft. K.Z. helped with SAMOSA data processing and feedback on computational work. V.R. and K.S.P. provided guidance and mentorship for entire work and thoroughly edited the manuscript. All co-authors reviewed and contributed to the final manuscript.

## Acknowledgements

We thank Geoffrey Fudenberg for input on adapting the Akita framework to SAMOSA data, Shuzhen Kuang for assistance with model interpretation tools and feedback on results, and members of the Ramani and Pollard lab for helpful discussions.

## References

1. Klemm, S. L., Shipony, Z. & Greenleaf, W. J. Chromatin accessibility and the regulatory epigenome. Nat Rev Genet 20, 207–220 (2019).

2. Zeitlinger, J. et al. Perspective on recent developments and challenges in regulatory and systems genomics. Bioinformatics Advances 5, vbaf106 (2024).

3. Sönmezer, C. et al. Molecular co-occupancy identifies transcription factor binding cooperativity in vivo. Mol. Cell 81, 255–267.e6 (2021).

4. Junion, G. et al. A Transcription Factor Collective Defines Cardiac Cell Fate and Reflects Lineage History. Cell 148, 473–486 (2012).

5. Tijssen, M. R. et al. Genome-wide Analysis of Simultaneous GATA1/2, RUNX1, FLI1, and SCL Binding in Megakaryocytes Identifies Hematopoietic Regulators. Developmental Cell 20, 597–609 (2011).

6. Buenrostro, J. D., Giresi, P. G., Zaba, L. C., Chang, H. Y. & Greenleaf, W. J. Transposition of native chromatin for fast and sensitive epigenomic profiling of open chromatin, DNA-binding proteins and nucleosome position. Nat. Methods 10, 1213–1218 (2013).

7. Thurman, R. E. et al. The accessible chromatin landscape of the human genome. Nature 489, 75–82 (2012).

8. Valouev, A. et al. Determinants of nucleosome organization in primary human cells. Nature 474, 516–520 (2011).

9. Krebs, A. R. et al. A practical guide to studying genome function using single-molecule genomics. Nat Rev Mol Cell Biol https://doi.org/10.1038/s41580-026-01017-4 (2026) doi:10.1038/s41580-026-01017-4.

10. Krebs, A. R. et al. Genome-wide single-molecule footprinting reveals high RNA polymerase II turnover at paused promoters. Mol. Cell 67, 411–422.e4 (2017).

11. Wang, Y. et al. Single-molecule long-read sequencing reveals the chromatin basis of gene expression. Genome Res. 29, 1329–1342 (2019).

12. Abdulhay, N. J. et al. Massively multiplex single-molecule oligonucleosome footprinting. Elife 9, (2020).

13. Shipony, Z. et al. Long-range single-molecule mapping of chromatin accessibility in eukaryotes. Nat. Methods 17, 319–327 (2020).

14. Stergachis, A. B., Debo, B. M., Haugen, E., Churchman, L. S. & Stamatoyannopoulos, J. A. Single-molecule regulatory architectures captured by chromatin fiber sequencing. Science 368, 1449–1454 (2020).

15. Lee, I. et al. Simultaneous profiling of chromatin accessibility and methylation on human cell lines with nanopore sequencing. Nat. Methods 17, 1191–1199 (2020).

16. Abdulhay, N. J. et al. Nucleosome density shapes kilobase-scale regulation by a mammalian chromatin remodeler. Nat. Struct. Mol. Biol. https://doi.org/10.1038/s41594-023-01093-6 (2023) doi:10.1038/s41594-023-01093-6.

17. Rao, S., Ahmad, K. & Ramachandran, S. Cooperative binding between distant transcription factors is a hallmark of active enhancers. Mol. Cell 81, 1651–1665.e4 (2021).

18. Swanson, E. G. et al. Mapping single-cell diploid chromatin fiber architectures using DAF-seq. Nat Biotechnol https://doi.org/10.1038/s41587-025-02914-3 (2025) doi:10.1038/s41587-025-02914-3.

19. Pampari, A. et al. ChromBPNet: bias factorized, base-resolution deep learning models of chromatin accessibility reveal cis-regulatory sequence syntax, transcription factor footprints and regulatory variants. bioRxivorg (2025) doi:10.1101/2024.12.25.630221.

20. Linder, J., Srivastava, D., Yuan, H., Agarwal, V. & Kelley, D. R. Predicting RNA-seq coverage from DNA sequence as a unifying model of gene regulation. Nat Genet 57, 949–961 (2025).

21. Avsec, Ž. et al. Advancing regulatory variant effect prediction with AlphaGenome. Nature 649, 1206–1218 (2026).

22. Avsec, Ž. et al. Base-resolution models of transcription-factor binding reveal soft motif syntax. Nat. Genet. 53, 354–366 (2021).

23. Ostrowski, M. S. et al. The single-molecule accessibility landscape of newly replicated mammalian chromatin. Cell https://doi.org/10.1016/j.cell.2024.10.039 (2024) doi:10.1016/j.cell.2024.10.039.

24. Yang, M. G. et al. Pervasive and programmed nucleosome distortion on single chromatin fibres. Nature https://doi.org/10.1038/s41586-026-10418-6 (2026) doi:10.1038/s41586-026-10418-6.

25. Rowley, M. J. & Corces, V. G. Organizational principles of 3D genome architecture. Nat Rev Genet 19, 789–800 (2018).

26. Hitz, B. C. et al. The ENCODE Uniform Analysis Pipelines. Preprint at http://biorxiv.org/lookup/doi/10.1101/2023.04.04.535623 (2023).

27. Davis, C. A. et al. The Encyclopedia of DNA elements (ENCODE): data portal update. Nucleic Acids Research 46, D794–D801 (2018).

28. The ENCODE Project Consortium. An integrated encyclopedia of DNA elements in the human genome. Nature 489, 57–74 (2012).

29. Martin, B. J. E. et al. Global identification of SWI/SNF targets reveals compensation by EP400. Cell 186, 5290–5307.e26 (2023).

30. Ramani, V., Qiu, R. & Shendure, J. High sensitivity profiling of chromatin structure by MNase-SSP. Cell Rep. 26, 2465–2476.e4 (2019).

31. ENCODE Project Consortium et al. Expanded encyclopaedias of DNA elements in the human and mouse genomes. Nature 583, 699–710 (2020).

32. Whalen, S., Schreiber, J., Noble, W. S. & Pollard, K. S. Navigating the pitfalls of applying machine learning in genomics. Nat Rev Genet 23, 169–181 (2022).

33. Fudenberg, G., Kelley, D. R. & Pollard, K. S. Predicting 3D genome folding from DNA sequence with Akita. Nat. Methods 17, 1111–1117 (2020).

34. Ovek Baydar, D. et al. JASPAR 2026: expansion of transcription factor binding profiles and integration of deep learning models. Nucleic Acids Research 54, D184–D193 (2026).

35. Kulakovskiy, I. V. et al. HOCOMOCO: towards a complete collection of transcription factor binding models for human and mouse via large-scale ChIP-Seq analysis. Nucleic Acids Research 46, D252–D259 (2018).

36. Wang, A. T. Fi-NeMo: Finding Neural network Motifs. Fi-NeMo: Finding Neural network Motifs https://github.com/austintwang/finemo_gpu (2024).

37. Shrikumar, A., Greenside, P. & Kundaje, A. Learning Important Features Through Propagating Activation Differences. https://doi.org/10.48550/ARXIV.1704.02685 (2017) doi:10.48550/ARXIV.1704.02685.

38. Shrikumar, A. et al. Technical Note on Transcription Factor Motif Discovery from Importance Scores (TF-MoDISco) version 0.5.6.5. Preprint at http://arxiv.org/abs/1811.00416 (2020).

39. Kuang, S. & Wang, L. Deep Learning of Sequence Patterns for CCCTC-Binding Factor-Mediated Chromatin Loop Formation. Journal of Computational Biology 28, 133–145 (2021).

40. Wen, Z., Huang, Z.-T., Zhang, R. & Peng, C. ZNF143 is a regulator of chromatin loop. Cell Biol Toxicol 34, 471–478 (2018).

41. Cerron-Alvan, L. M. et al. Wt1 splice isoforms bias lineage programmes during formative pluripotency. Preprint at http://biorxiv.org/lookup/doi/10.64898/2026.04.06.713568 (2026).

42. Michael, A. K. et al. Mechanisms of OCT4-SOX2 motif readout on nucleosomes. Science 368, 1460–1465 (2020).

43. Owens, N. et al. CTCF confers local nucleosome resiliency after DNA replication and during mitosis. Elife 8, (2019).

44. Ceribelli, M. et al. The Histone-Like NF-Y Is a Bifunctional Transcription Factor. Molecular and Cellular Biology 28, 2047–2058 (2008).

45. Iliadou, M., Haro, E. & Rada-Iglesias, A. Mechanisms and functional implications of long-range enhancer-dependent gene regulation. Nat Genet 58, 2104–2117 (2026).

46. Jolma, A. et al. An expanded codebook of human transcription factor DNA-binding specificity. Nature 657, 275–283 (2026).

47. Bell, J. S. K. & Vertino, P. M. Orphan CpG islands define a novel class of highly active enhancers. Epigenetics 12, 449–464 (2017).

48. Yang, J., Corsello, T. R. & Ma, Y. Stem Cell Gene SALL4 Suppresses Transcription through Recruitment of DNA Methyltransferases. Journal of Biological Chemistry 287, 1996–2005 (2012).

49. Besson, J.-M. et al. Selective disruption of DNMT1/ELK1 interactions induces DGKI re-expression and promotes temozolomide sensitivity of MGMTmethylated/DGKImethylated glioblastoma. Clin Epigenet 17, 146 (2025).

50. Li (李怡灵), Y. et al. Transcriptional repression by TGIF2 coordinates neurogenic priming and neural stem cell maintenance. Sci. Adv. 12, eaea9974 (2026).

51. Bernardini, A. & Mantovani, R. Q-rich activation domains: flexible ‘rulers’ for transcription start site selection? Trends in Genetics 41, 275–285 (2025).

52. Domcke, S. et al. Competition between DNA methylation and transcription factors determines binding of NRF1. Nature 528, 575–579 (2015).

53. Brandeis, M. et al. Spl elements protect a CpG island from de novo methylation. Nature 371, 435–438 (1994).

54. Dolfini, D., Gatta, R. & Mantovani, R. NF-Y and the transcriptional activation of CCAAT promoters. Critical Reviews in Biochemistry and Molecular Biology 47, 29–49 (2012).

55. Hartl, D. et al. CG dinucleotides enhance promoter activity independent of DNA methylation. Genome Res. 29, 554–563 (2019).

56. Kaluscha, S. et al. Evidence that direct inhibition of transcription factor binding is the prevailing mode of gene and repeat repression by DNA methylation. Nat Genet 54, 1895–1906 (2022).

57. Parker, J. B., Yin, H., Vinckevicius, A. & Chakravarti, D. Host Cell Factor-1 Recruitment to E2F-Bound and Cell-Cycle-Control Genes Is Mediated by THAP11 and ZNF143. Cell Reports 9, 967–982 (2014).

58. Grand, R. S. et al. BANP opens chromatin and activates CpG-island-regulated genes. Nature 596, 133–137 (2021).

59. Gill, G., Pascal, E., Tseng, Z. H. & Tjian, R. A glutamine-rich hydrophobic patch in transcription factor Sp1 contacts the dTAFII110 component of the Drosophila TFIID complex and mediates transcriptional activation. Proc. Natl. Acad. Sci. U.S.A. 91, 192–196 (1994).

60. Ferreri, K., Gill, G. & Montminy, M. The cAMP-regulated transcription factor CREB interactswith a component of the TFIID complex. Proc. Natl. Acad. Sci. U.S.A. 91, 1210–1213 (1994).

61. Coustry, F., Sinha, S., Maity, S. N. & Crombrugghe, B. D. The two activation domains of the CCAAT-binding factor CBF interact with the dTAFII110 component of the Drosophila TFIID complex. Biochemical Journal 331, 291–297 (1998).

62. Sun, X., Zhang, J. & Cao, C. CTCF and Its Partners: Shaper of 3D Genome during Development. Genes 13, 1383 (2022).

63. Julian, L. M. et al. Tissue-specific targeting of cell fate regulatory genes by E2f factors. Cell Death Differ 23, 565–575 (2016).

64. Malinen, M. et al. Distinct HDACs regulate the transcriptional response of human cyclin-dependent kinase inhibitor genes to trichostatin A and 1α,25-dihydroxyvitamin D 3. Nucleic Acids Research 36, 121–132 (2008).

65. Lodewijk, G. A. et al. E2F1 induces a G0-G1 reentry transcriptional program without changing chromatin accessibility. Preprint at http://biorxiv.org/lookup/doi/10.1101/2025.09.23.678145 (2025).

66. Huang, H. et al. CTCF mediates dosage- and sequence-context-dependent transcriptional insulation by forming local chromatin domains. Nat Genet 53, 1064–1074 (2021).

67. Fleming, J. D. et al. NF-Y coassociates with FOS at promoters, enhancers, repetitive elements, and inactive chromatin regions, and is stereo-positioned with growth-controlling transcription factors. Genome Res. 23, 1195–1209 (2013).

68. Zhou, J. & Troyanskaya, O. G. Predicting effects of noncoding variants with deep learning-based sequence model. Nat. Methods 12, 931–934 (2015).

69. Kelley, D. R. et al. Sequential regulatory activity prediction across chromosomes with convolutional neural networks. Genome Res. 28, 739–750 (2018).

70. Avsec, Ž. et al. Effective gene expression prediction from sequence by integrating long-range interactions. Nat. Methods 18, 1196–1203 (2021).

71. Chen, K. M., Wong, A. K., Troyanskaya, O. G. & Zhou, J. A sequence-based global map of regulatory activity for deciphering human genetics. Nat. Genet. 54, 940–949 (2022).

72. Pachano, T. et al. Orphan CpG islands amplify poised enhancer regulatory activity and determine target gene responsiveness. Nat Genet 53, 1036–1049 (2021).

73. Wang, J. et al. Sequence features and chromatin structure around the genomic regions bound by 119 human transcription factors. Genome Res. 22, 1798–1812 (2012).

74. Boulanger, M. et al. Molecular co-accessibility identifies coordinated regulation between distant cis-regulatory elements. Molecular Cell 86, 3394–3410.e8 (2026).

75. Zhang, K., Maristany, M. J., Huertas, J., Collepardo-Guevara, R. & Ramani, V. Single-molecule nucleosome spacing coordinates chromatin fiber interactions. bioRxivorg (2026) doi:10.64898/2026.06.02.729033.

76. Reiff, S. B. et al. The 4D Nucleome Data Portal as a resource for searching and visualizing curated nucleomics data. Nat Commun 13, 2365 (2022).

77. Murphy, D. et al. 3D Enhancer–promoter networks provide predictive features for gene expression and coregulation in early embryonic lineages. Nat Struct Mol Biol 31, 125–140 (2024).

78. Gupta, S., Stamatoyannopoulos, J. A., Bailey, T. L. & Noble, W. S. Quantifying similarity between motifs. Genome Biol. 8, R24 (2007).

79. Khan, A., Riudavets Puig, R., Boddie, P. & Mathelier, A. BiasAway: command-line and web server to generate nucleotide composition-matched DNA background sequences. Bioinformatics 37, 1607–1609 (2021).

80. Illingworth, R. S. et al. Orphan CpG Islands Identify Numerous Conserved Promoters in the Mammalian Genome. PLoS Genet 6, e1001134 (2010).

81. Gardiner-Garden, M. & Frommer, M. CpG islands in vertebrate genomes. J Mol Biol 196, 261–282 (1987).

