## Supplemental Material for "ChAI-Net learns a regulatory code for chromatin co-accessibility"

### Extended Data

Extended Data Figure 1
Extended Data Figure 2
Extended Data Figure 3
Extended Data Figure 4
Extended Data Figure 5
Extended Data Figure 6
Extended Data Figure 7
Extended Data Figure 8
Extended Data Figure 9

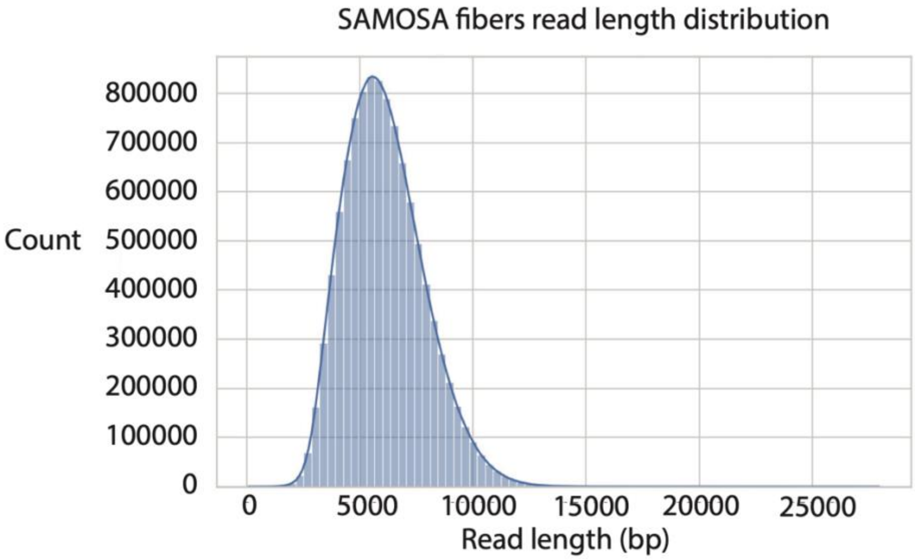


**Extended Data 1. Distribution of read lengths of SAMOSA fibers used for model training.** Histogram of the number of reads across the range of read lengths.

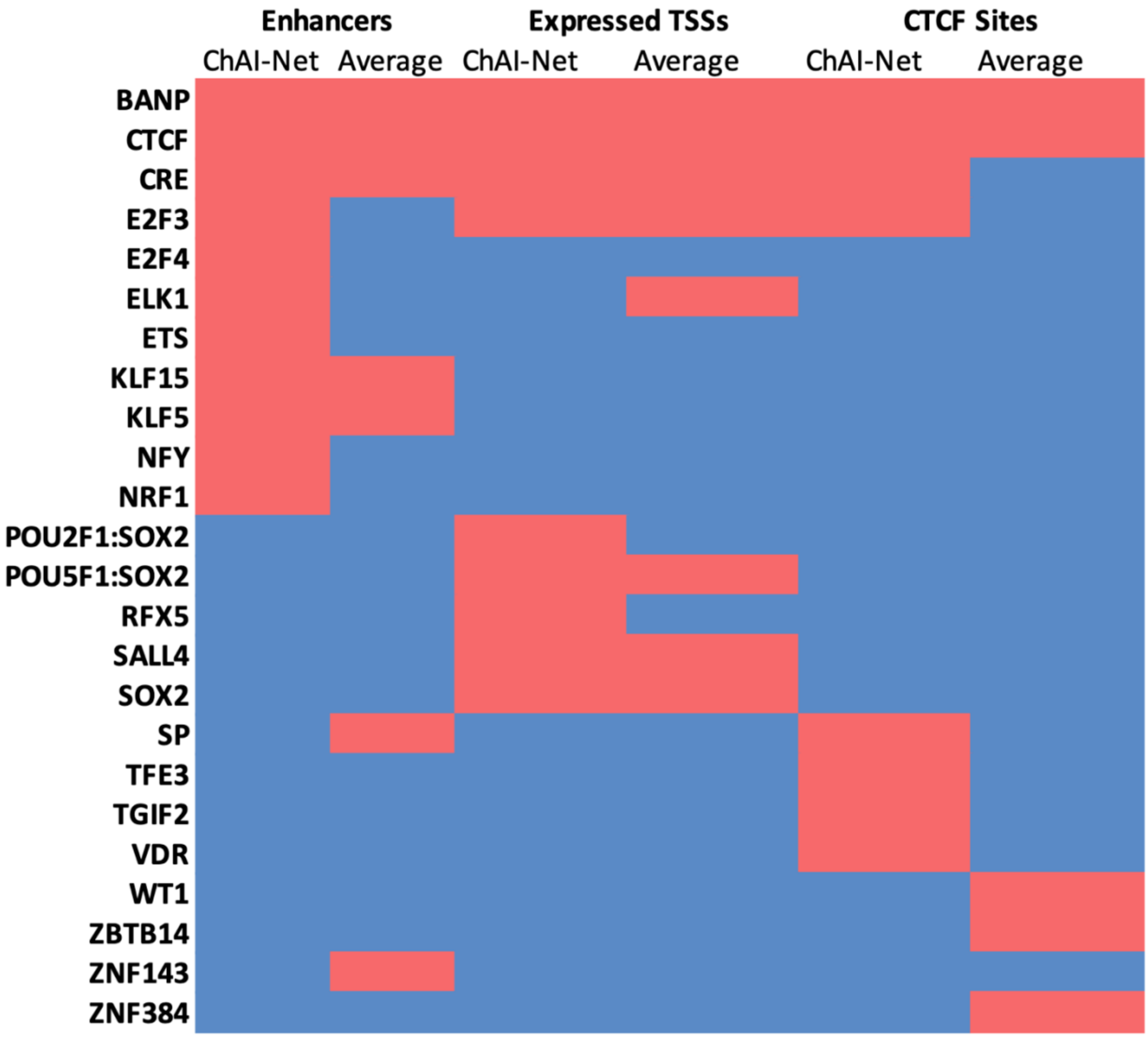


**Extended Data 2. Comparison of TF PWM matching *de novo* motifs learned by ChAI-Net and the average accessibility model across regulatory regions.** Rows: TFs whose PWM matches a CWM from ChAI-Net and/or the average accessibility model. Columns: Source of CWM (regulatory region type, model). Values: Red = TF's PWM matches a CWM in this context indicating it is important for predicting co-accesibility; Blue: No match in this context. If two CWMs match the same TF, these are collapsed. For CTCF, we also collapsed multiple PWMs: CTCF-forward, CTCF-reverse complement, CTCF-extended, CTCFL. The full set of CWMs and PWMs can be seen in **Extended Data Figure 3** (ChAI-Net) or **Extended Data Figure 5** (average accessibility model).

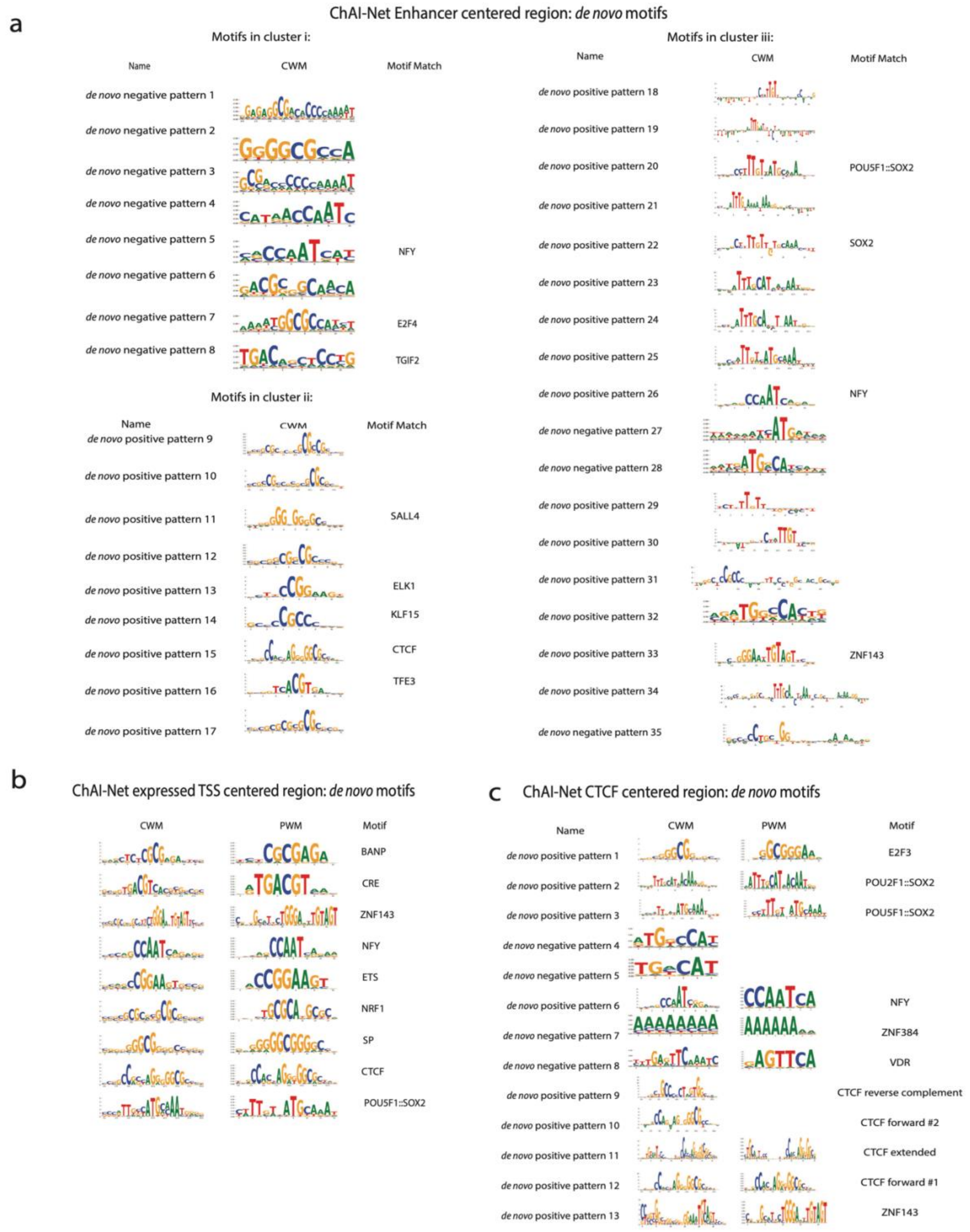


**Extended Data 3. ChAI-Net discovered *de novo* motifs across regulatory regions. a.** ChAI-Net *de novo* motifs learned from mESC enhancers ordered and grouped by the co-occurring motif clusters defined in **Fig. 4a**. Left: contribution weight matrix (CWM) name, Middle: CWM values, Right: name of position weight matrix (PWM) matching transcription factor (TF), if any. **b.** ChAI-Net *de novo* motifs learned from mESC expressed TSSs. Left: CWM, Middle: matching PWM, Right: TF name. **c.** ChAI-Net *de novo* motifs learned from mESC CTCF sites. Left: CWM name, Second from left: CWM, Third from left: PWM, Right: TF name.

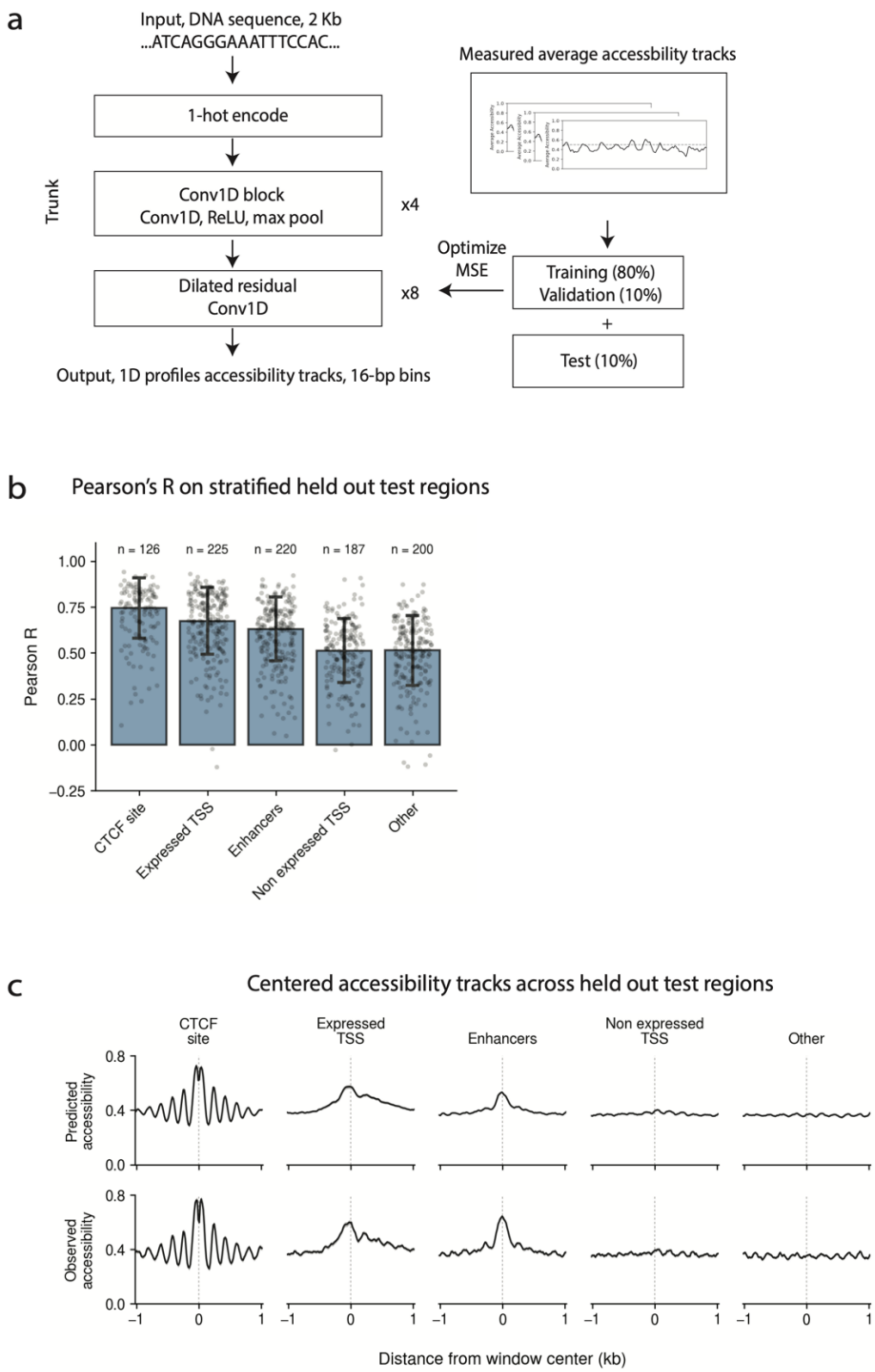


**Extended Data 4. Modeling SAMOSA average accessibility from DNA sequence. a.** Neural network architecture for modeling SAMOSA average accessibility from DNA sequence. The input is 2,048 bp of 1-hot encoded DNA. This is converted using a number of convolution and downsampling layers and a linear activation to predict a track of average accessibility at a resolution of 16-bp per bin. This model was trained using the same 80/10/10 training/validation/test splits used to train ChAI-Net (**Fig. 2A**). **b.** Average accessibility model held-out test window performance as Pearson's R stratified by windows overlapping regulatory elements (CTCF sites, expressed TSSs, enhancers, non-expressed TSSs) and other windows that do not overlap these regulatory annotations. **c.** Predicted and observed SAMOSA accessibility tracks averaged across test windows overlapping each type of regulatory element.

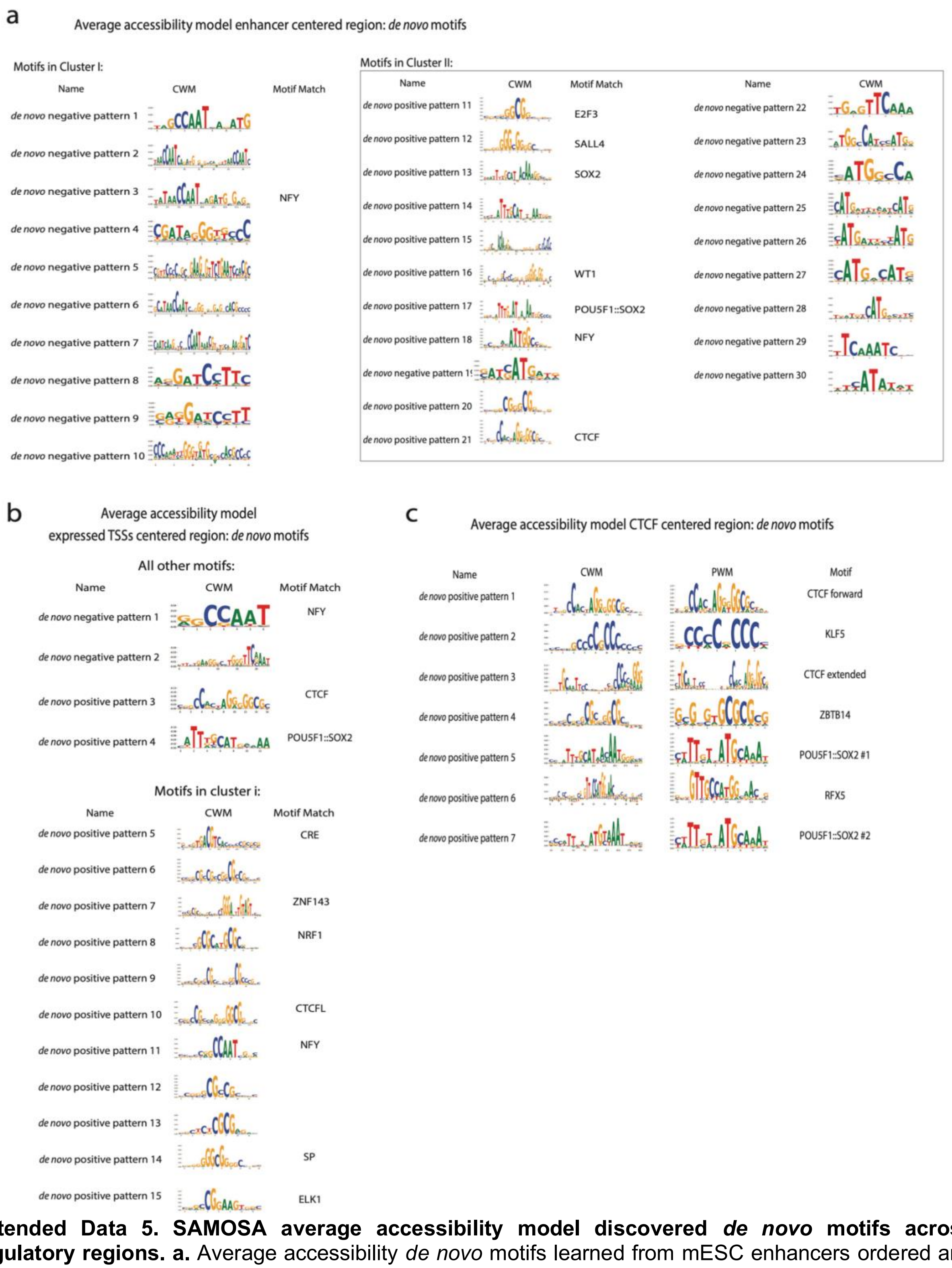


**Extended Data 5. SAMOSA average accessibility model discovered *de novo* motifs across regulatory regions. a.** Average accessibility *de novo* motifs learned from mESC enhancers ordered and grouped by the co-occurring motif clusters defined in **Fig. 4a**. Left: contribution weight matrix (CWM) name, Middle: CWM values, Right: name of position weight matrix (PWM) matching transcription factor (TF), if any. **b.** Average accessibility *de novo* motifs learned from mESC expressed TSSs. Left: CWM name, Middle: CWM, Right: TF name. **c.** Average accessibility *de novo* motifs learned from mESC CTCF sites. Left: CWM name, Second from left: CWM, Third from left: PWM, Right: TF name.

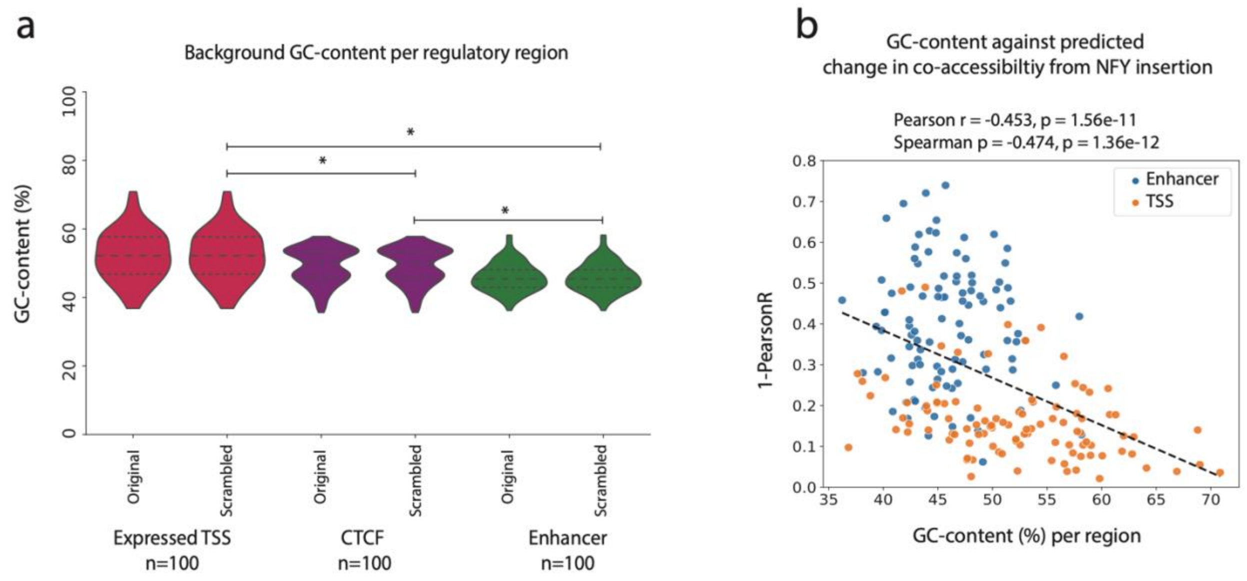


**Extended Data 6. GC-content in ChAI-Net co-accessibility predictions. a.** GC-content distributions across windows overlapping expressed TSSs, CTCF sites, and enhancers (Original) versus matched background sequences used for *in silico* experiments in **Fig. 3b** (Scrambled; n=100 per regulatory element type) for background DNA sequences (n=100). Star symbols indicate statistical difference (MWU $p < 0.05$). **b.** GC-content versus predicted effect of inserting the NFY CTCF site CWM consensus sequence into the 100 background DNA sequences for enhancers or expressed TSSs. Pearson and Spearman correlation are calculated using all 200 data points.

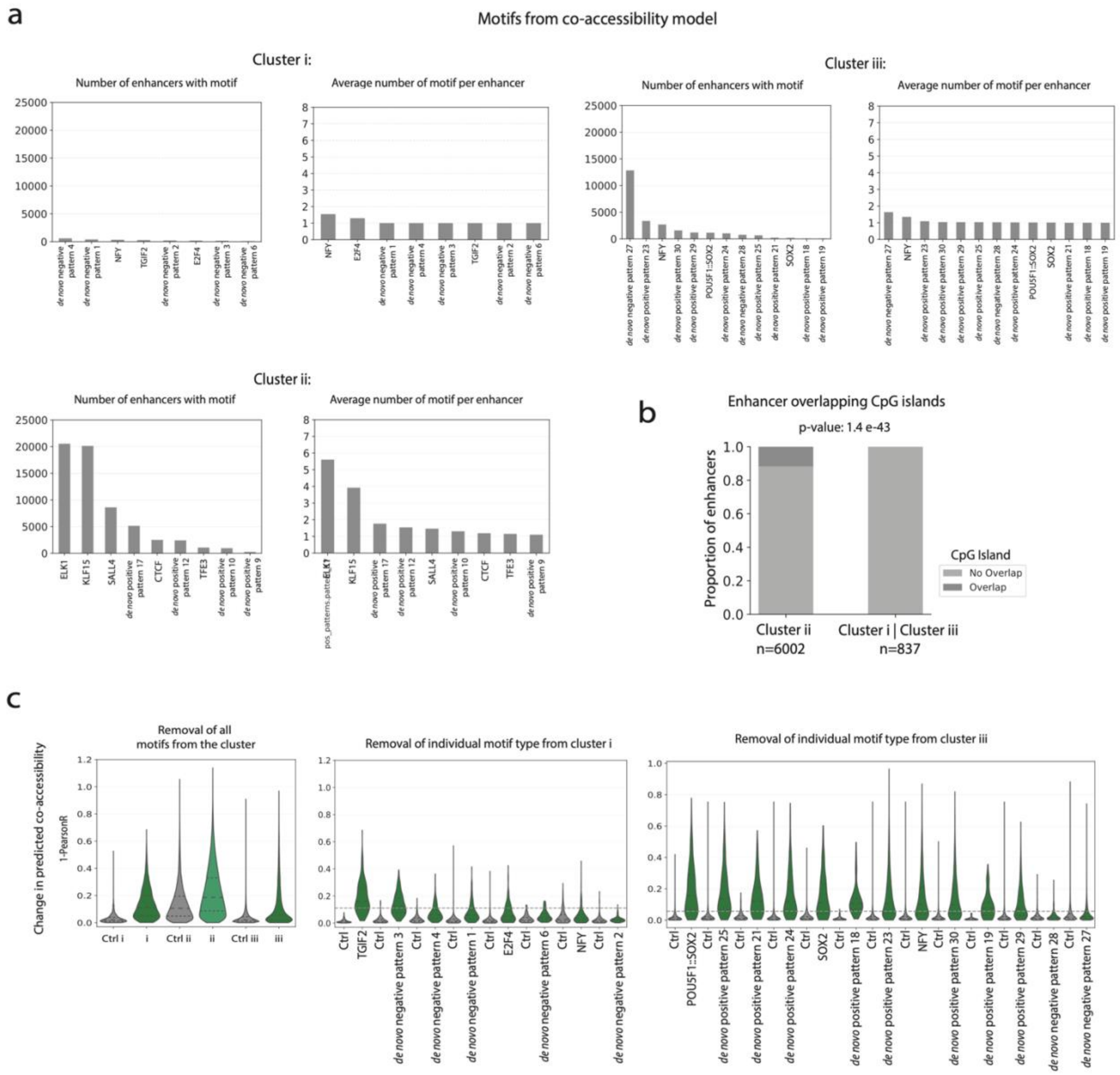


**Extended Data 7. Characterization of ChAI-Net discovered *de novo* motifs from enhancers. a.** Summary of the total number of enhancers with at least one instance of *de novo* motifs (left histograms in each set) and average counts of each motif per enhancer (right histograms), stratified by membership in ChAI-Net clusters as defined in **Fig. 4a**. **b.** Proportion of enhancers in cluster ii versus the combination of clusters i and iii that overlap CpG islands (CGIs). A one-sided Fisher's exact test of the null hypothesis that cluster ii does not have a higher rate of CGI overlap is reported (p=1.4e=43) with n denoting the sample size per group. **c.** Change in predicted co-accessibility upon motif ablation. Left: Ablating all motif instances from each cluster (shades of green) altered co-accessibility more than ablating matched controls (grey) (MWU test BH-adjusted $p < 0.05$ for all three clusters). Clusters varied in the number of enhancer windows: cluster i (n=1,570), cluster ii (n=22,823), cluster iii (n=16,335). This panel is identical to **Fig. 4f**, repeated here for comparison with the middle and right panels. Middle: Ablating instances of individual motifs from cluster i (green) altered co-accessibility more than their matched controls (all MWU test BH-adjusted $p < 0.05$) but generally less or similar to the combination of all motifs (median cluster i: dashed line), with the exception of TGIF2. Right: Ablating instances of individual motifs from cluster iii (green) altered co-accessibility more than their matched controls (all MWU test BH-adjusted $p < 0.05$) and in some cases similar to the combination of all motifs (median cluster iii: dashed line).

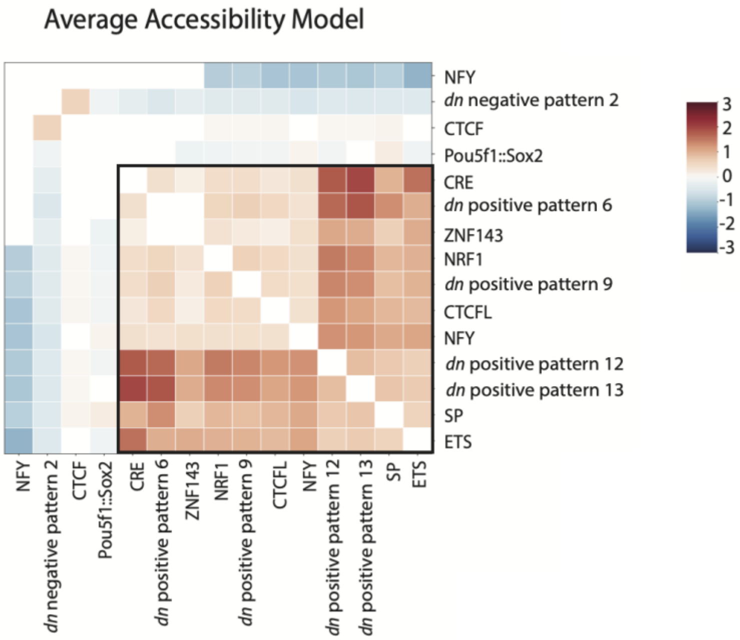


**Extended Data 8. Co-occurrence of *de novo* motifs learned from expressed TSSs by the average accessibility model.** Co-occurring *de novo* motifs, with CWM-PWM matching TFs labeled. Significantly (Fisher's exact test BH-adjusted $p < 0.05$) enriched (red; log2 odds ratio) or depleted (blue; log2 odds ratio) motif pairs are clustered based on their co-occurrence patterns across TSS centered sites. Box: a cluster of motifs enriched for co-occurring in the same expressed TSSs.

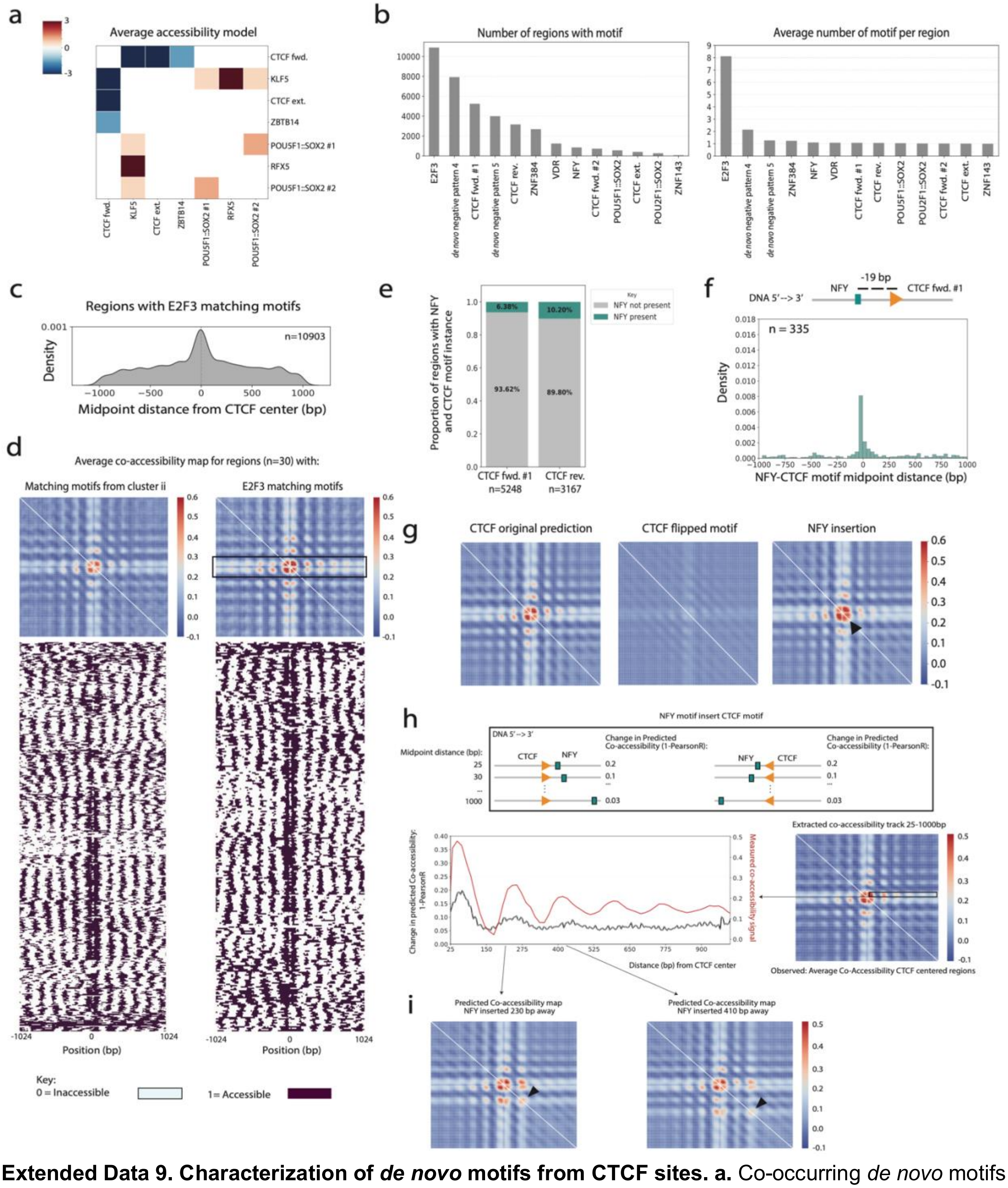


**Extended Data 9. Characterization of *de novo* motifs from CTCF sites. a.** Co-occurring *de novo* motifs learned by the average accessibility mode, with TF PWM matches labeled. Significantly (Fisher's exact test BH-adjusted $p < 0.05$) enriched (red; log2 odds ratio) or depleted (blue; log2 odds ratio) motif pairs are clustered based on their co-occurrence patterns across CTCF sites. **b.** Summary of the total number of CTCF sites with at least one instance of *de novo* motifs from ChAI-Net (left) and average counts of each motif per CTCF site (right). **c.** Distribution of midpoint distances between instances of ChAI-Net E2F3 motifs and CTCF motifs (n=10,903 CTCF sites). **d.** Observed co-accessibility map averages for windows most sensitive to *in silico* ablation of all cluster ii motifs (left; defined in **Fig. 6b**) or all E2F3 motifs (right). Shown below are the SAMOSA fibers from all windows contributing to each average. Black box: higher co-accessibility at sites flanking the CTCF motif. **e.** Proportion of CTCF sites with NFY motifs when CTCF is in the forward orientation (left) versus reverse complement orientation (right). **f.** Distribution of motif-to-motif midpoint distances for CTCF-forward and NFY instances within the same CTCF site window. The most common distance (~19 bp) is labeled in the cartoon above. **g.** Co-accessibility predictions averaged over 658 windows with CTCF and NFY motifs in their original orientations (left), with CTCF motif orientations flipped (middle), and with NFY inserted in front of CTCF (right). Black arrow: wider co-accessible region proximal to the NFY insertion. Both CTCF-forward and CTCF-reverse complement motifs are included, but CTCF-reverse complement examples are flipped so that all windows have the same CTCF orientation. **h.** Schematic of NFY insertion experiments. Top: Motifs were inserted at 5-bp intervals from 25 to 1000 bp from the CTCF motif midpoint. Co-accessibility maps were predicted before and after the insertion, with the change quantified using 1-Pearson's correlation (1-*r*). Bottom: median change in predicted co-accessibility (black; 1-*r*) for all modified sequences as a function of motif insertion distance from the CTCF motif midpoint, alongside observed co-accessibility with the CTCF bin (red). **i.** Representative predicted co-accessibility maps from *in silico* NFY insertion experiments (**h**) showing NFY CWM consensus sequences inserted 230 bp (left map) and 410 bp (right map) away from the central CTCF motif. Black arrows: co-accessible sites.
